# Cancer cells acquire cysteine directly from a reduced tumor microenvironment

**DOI:** 10.64898/2026.10.25.761647

**Authors:** Feroza K. Choudhury, Basma A. Yasseen, Paloma González-Sánchez, Meredith Long, Samantha Caldwell, Pritin N. Soni, Suehelay Acevedo-Acevedo, Yun Huang, Elsa R. Flores, Chang Jiang, Nathan P. Ward, Gina M. DeNicola

**Author notes:** Contributed Equally (Feroza K. Choudhury, Basma A. Yasseen).

## Abstract

The metabolic environment of a tumor is shaped by the nutrients available in its interstitial fluid, yet the redox composition of that fluid is poorly defined. Measuring sulfur metabolites in the interstitial fluid of tumors, matched healthy organs, and serum across three mouse models of cancer, we find that the tumor microenvironment (TME) is a chemically reduced compartment in which free cysteine predominates over its oxidized form, cystine. This is especially pronounced in the lung TME, where cystine is nearly absent. We further show that lung cancer cells take up cysteine and cystine with comparable efficiency and proliferate on cysteine alone, and that the abundant glutathione in the TME helps keep cysteine reduced through disulfide exchange. In a medium formulated to match the sulfur composition of the lung tumor interstitial fluid, cells become independent of cystine import through xCT. Cells cultured in this medium were resistant to glutaminase inhibitors, oxidants and ferroptosis inducers, and better maintained NADPH levels under oxidative challenge. These findings suggest that standard culture conditions overestimate the dependence of cancer cells on cystine uptake and their vulnerability to oxidative stress-targeting therapies.

## Main

Cysteine is the limiting substrate for glutathione (GSH) synthesis and a required precursor for coenzyme A, iron–sulfur clusters, and protein synthesis^1^. Much of what is known about how cancer cells acquire it comes from cell culture, where cysteine is supplied almost exclusively as cystine because free cysteine autoxidizes under the oxygen tension, pH, and trace metal content of standard media^2^. Under these conditions, cancer cells depend on the cystine/glutamate antiporter xCT (SLC7A11), and the imported cystine must be reduced at the expense of NADPH before it can be used^3^. This model underlies much of what is inferred about ferroptosis sensitivity, glutamate export, and disulfide stress^4–6^. Systemic depletion of cyst(e)ine restricts tumor growth^7,8^, and isotope tracing places the origin of tumor cysteine in the circulation^9^, where it exists predominantly as the oxidized disulfide. Interstitial fluid (IF) profiling has shown, however, that the composition of tissue microenvironments differs substantially from that of plasma^10,11^. However, because thiols oxidize during IF collection and extraction, the relative abundance of cysteine and cystine in the tumor microenvironment (TME) has not been accurately captured.

Here we quantified sulfur amino acid and glutathione metabolites in tumor IF, matched healthy organ IF, and serum with their native redox state preserved. Across mouse models of lung, liver, and pancreatic cancer, the TME was a chemically reduced compartment dominated by cysteine and GSH, whereas serum and healthy organ IF contained more abundant cystine and GSSG. GSH sustained this state by reducing cystine through a cysteinyl-glutathione intermediate, making cysteine the predominant bioavailable species. The reduced state was not cancer cell– intrinsic as it was lost at a non-native tissue site and traced in part to tumor-associated macrophages. In medium mimicking lung tumor IF, cancer cells became independent of xCT, and resistant to glutaminase inhibition and oxidative insults. Widely studied cystine-centric dependencies are therefore largely a property of culture, and the reduced TME determines which redox pathways are required in vivo.

### The tumor microenvironment is chemically reduced across cancer types

To assess sulfur metabolite availability in the tumor microenvironment, we analyzed absolute concentrations of metabolites in cysteine/glutathione redox metabolism in the interstitial fluid (IF) of tumors and matched healthy organ, alongside serum, using three genetically engineered mouse models (GEMMs) that develop organ-specific tumors: liver (Myc; Trp53^-/-^), lung (Kras^G12D/+^; Trp53^fl/fl^), and pancreas (Kras^G12D/+^; Trp53^fl/fl^; p48-cre) (Fig. 1a). Consistent with prior reports that tumor interstitial fluid is compositionally distinct from plasma^11^, tumor IF differed from matched serum across most measured metabolites in all three models (Fig. 1b, Supplementary Table 1). Compared to IF from the corresponding healthy organ, however, the differences were far more restricted, with the largest fold changes confined to metabolites of sulfur amino acid and glutathione metabolism (Fig. 1b). This signature was most pronounced in cysteine and its oxidized form cystine. In LUAD, tumor IF contained 36.2 µM cysteine, in sharp contrast to the negligible levels in serum or healthy lung (Fig. 1c), while cystine showed the opposite pattern, with less than 2 µM in tumor IF versus ∼15 µM in serum and lung (Fig. 1d). Both reduced (1.1mM) and oxidized (0.99mM) glutathione were markedly elevated in LUAD tumor IF (Fig. 1e,f). PDAC showed a similar cysteine/cystine relationship relative to serum and pancreas, whereas HCC and healthy liver contained predominantly the reduced form. Total glutathione differed across models: healthy liver reached millimolar concentrations, consistent with its central role in glutathione biosynthesis; HCC tumor IF was dominated by reduced glutathione; and PDAC had lower total glutathione than the other two tumor types, although significantly more than normal pancreas. Together, the data suggests TME across three distinct cancer types is a chemically reduced compartment enriched in cysteine in contrast to serum or healthy organs.

**Figure 1:**
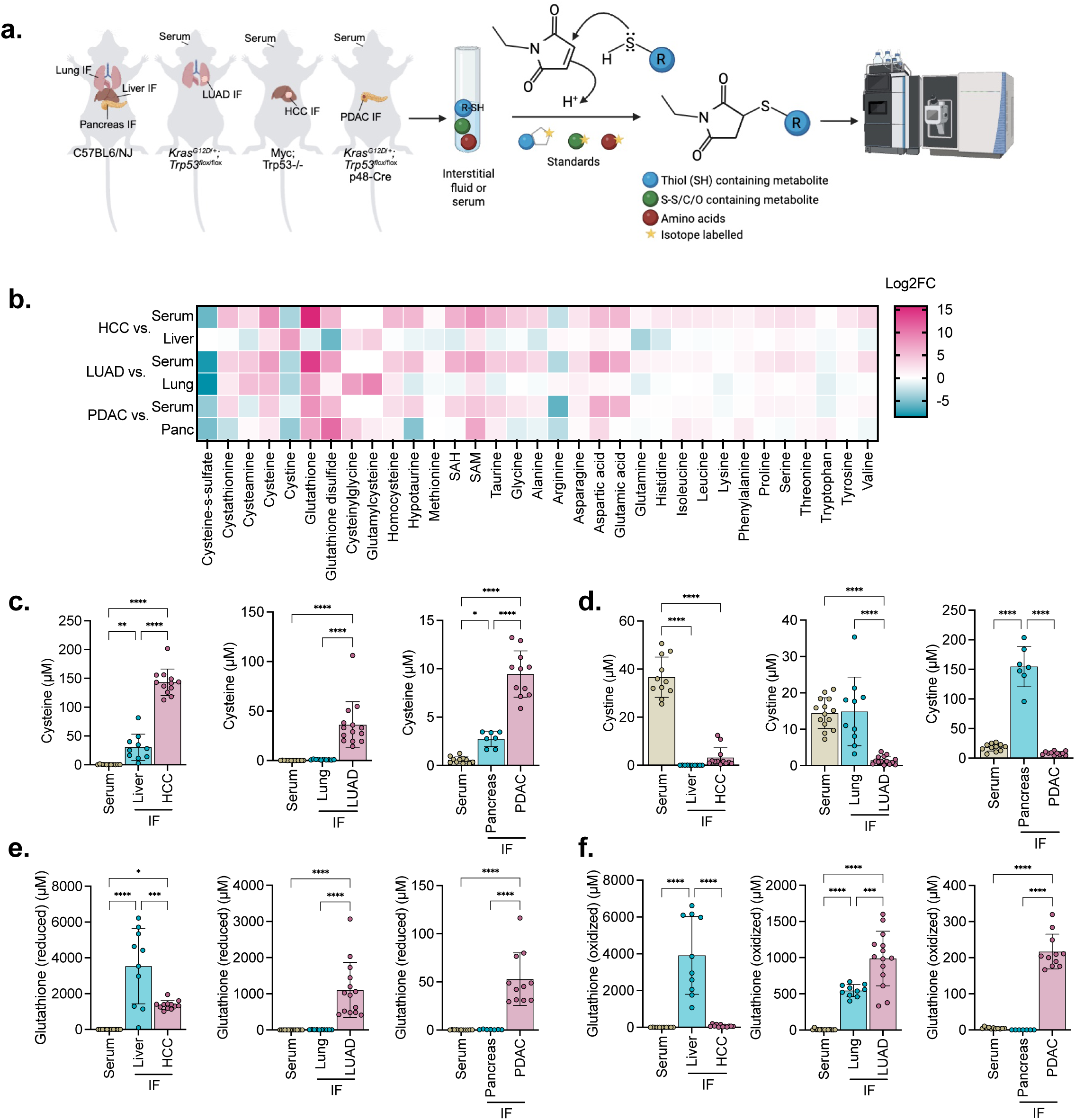
The tumor microenvironment is chemically reduced across cancer types. **(a)** Workflow for metabolite quantification from interstitial fluid. **(b)** Heatmap of average log2 fold change (FC) in metabolite concentration in tumor interstitial fluid versus matched serum or healthy organ interstitial fluid. **(c-e)** Concentration of cysteine (**c**), cystine (**d**), and reduced (**e**) and oxidized (**f**) glutathione in interstitial fluid of the indicated healthy organ and tumor, and serum from matched tumor-bearing mice. Data are mean ± s.d.; Liver: n = 12 (HCC), 10 (liver), 11 (serum). Lung: n = 14 (LUAD), 10 (lung), 14 (serum). Pancreas: n = 11 (PDAC), 7 (pancreas), 11 (serum). Statistical analysis was performed by one-way ANOVA with Tukey’s multiple comparisons test.

### Glutathione keeps cysteine in its reduced form to confer cystine independence

Free cysteine is chemically unstable and spontaneously oxidizes to cystine, raising the question of how the TME sustains cysteine in its reduced form. Indeed, incubating 100μM cysteine in cystine-free RPMI showed a half-life of approximately 10 hours, with cysteine loss mirrored by cystine accumulation (Fig. 2a). Critically, the oxidation rate increased ten-fold upon addition of dialyzed FBS (Fig. 2a). Because transition metals, particularly copper, catalyze thiol oxidation, we measured copper by ICP-MS and found it significantly elevated in serum-containing medium (Fig. S1), implicating serum-derived copper as a driver of accelerated oxidation.

**Figure 2:**
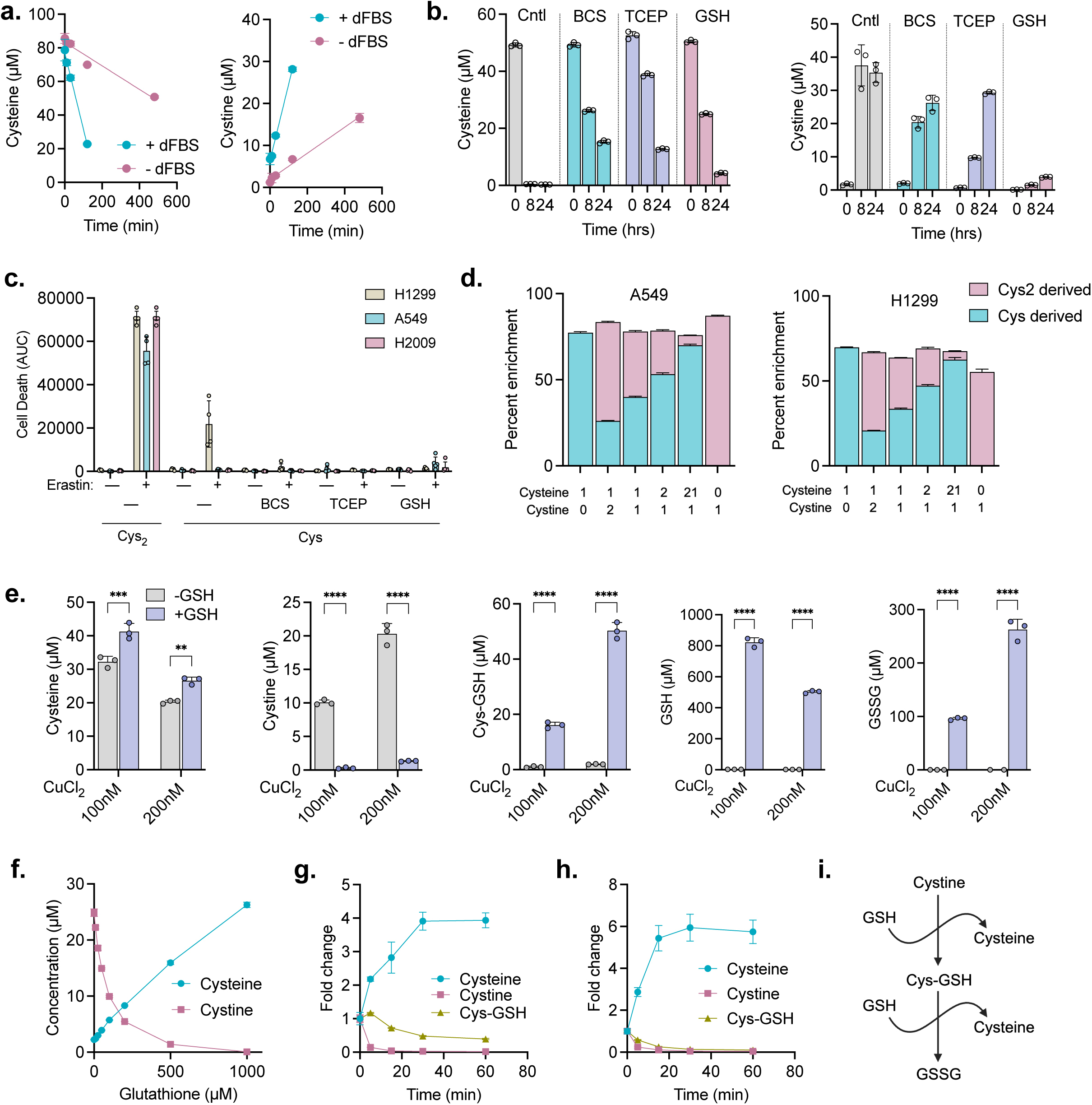
Glutathione is critical to maintain the cysteine redox state. **(a)** Cysteine and cystine levels over time in cell-free medium supplemented with 100µM cysteine, with or without 10% dialyzed FBS (dFBS) after 0, 10, 30, 120 and 480 min of incubation. **(b)** Cysteine and cystine levels over time (0, 8 and 24 h) in cell-free RPMI medium +10% dFBS supplemented with vehicle, 20 µM BCS, 100 µM TCEP, or 1mM GSH. **(c)** Quantification of Erastin-induced cell death in H1299, A549, and H2009 cells cultured in cystine-containing medium or cysteine-containing medium supplemented with 20 µM BCS, 100 µM TCEP, or 1mM GSH. Cell death was quantified as area under the curve of SYTOX Green-positive cells normalized to cell confluence over 72 h of live-cell imaging.**(d)** Percent enrichment of ^13^C and ^34^S — derived from [U-^13^C]-cysteine and ^34^S-cystine, respectively, in A549 and H1299 cells following a 30min incubation with [U-^13^C]cysteine and ^34^S-cystine at the indicated ratios, expressed in cysteine equivalents (44 µM total cysteine equivalents; cystine concentrations were calculated as half the stated equivalent value, reflecting its two cysteine residues per molecule).**(e)** Cysteine, cystine, cysteinyl-glutathione (Cys-GSH), glutathione (GSH) and glutathione disulfide (GSSG) levels following incubation of 100µM cysteine, with or without 1mM GSH, in RPMI containing 100nM or 200nM CuCl_2_ for 1 h. Statistical analysis was performed by two-way ANOVA with Sidak’s multiple comparison test. **(f)** Cysteine and cystine levels following incubation of 100µM cysteine with the indicated concentration of GSH in RPMI + 10% dFBS for 4 h. **(g)** Relative cysteine, cystine and Cys-GSH levels following incubation of 100µM cystine with 1mM GSH in RPMI + 10% dFBS for 5, 15, 30 and 60 min. **(h)** Relative cysteine, cystine and Cys-GSH levels following incubation of 100µM Cys-GSH with 1mM GSH in RPMI + 10% dFBS for 5, 15, 30 and 60 min. All data are mean ± s.d. (**i**) Schematic of the disulfide exchange reaction between cystine and glutathione (GSH) where cysteinyl-glutathione (Cys-GSH) is produced in the first step, which further reacts with GSH to produce oxidized glutathione (GSSG) and liberates cysteine in both steps. All data are mean ± s.d. n = 3 (**a,b,d-h**) or n = 4 (**c**) biological replicates, with at least 2 (**a,b,e-h**) or 3 (**c,d**) independent experiments.

We tested three strategies to stabilize cysteine in its reduced form: the copper chelator bathocuproine disulfonic acid (BCS); the cell impermeable, non-thiol reducing agent tris(2-carboxyethyl)phosphine (TCEP); and glutathione, the physiological reductant found in tumor IF. Each preserved cysteine while limiting cystine accumulation; notably, GSH did so at the concentration we measured in lung tumor IF (Fig. 2b). We then tested whether stabilizing extracellular cysteine impacts xCT dependence. H1299, A549, and H2009 cells were cultured in either cystine-containing medium or cysteine-containing medium supplemented with BCS, TCEP, or GSH, and challenged with erastin. Addition of cysteine alone or with its stabilizers significantly reduced erastin-induced cell death compared to cystine-containing medium, decreasing the area under the cell-death curve across all three conditions and cell lines (Fig. 2c, Fig. S2a).

To determine whether NSCLC cells can uptake cysteine directly, we cultured A549 and H1299 cells with mixtures of [U-^13^C]-cysteine and ^34^S_2_-cystine at varying cysteine-equivalent ratios for 30 minutes, using ^13^C-and ^34^S-labeling to trace the contribution of each extracellular source to intracellular cysteine. The isotopic enrichment of intracellular cysteine closely tracked with the extracellular cysteine-equivalent ratio of cysteine to cystine (e.g. a 1:1 cysteine-equivalent ratio yielded 50:50 intracellular labeling, Fig. 2d), indicating that lung cancer cells take up cysteine and cystine with comparable efficiency on a per-cysteine-equivalent basis. Collectively, these results indicate that cysteine uptake can bypass xCT dependence, although this protective effect may be limited by cysteine oxidation in culture.

We next considered γ-glutamyl transpeptidase (GGT), which catabolizes GSH extracellularly and could contribute to the bioavailable cysteine pool^12^. However, two lines of evidence argue against this route in lung cancer. First, the GGT product cysteinylglycine was abundant in HCC IF but was present at only 3μM in LUAD IF (Fig. S2b), indicating limited extracellular GSH catabolism in the lung TME. Second, we tested the GGT dependence directly by treating H1299 cells with the GGT inhibitor GGsTOP and assessed GSH-mediated protection against erastin in cysteine-containing medium. GGsTOP did not diminish this protection (Fig. S2c), indicating that GSH does not act as a cysteine donor in this setting. Moreover, in tracing experiments when we labelled the H1299 cells with [U-^13^C]-Cysteine, in presence of unlabeled glutathione, labelling of cysteine was reduced by only ∼3%, indicating very little contribution of glutathione catabolism (Fig. S2d). GSH sequesters copper intracellularly to limit copper-mediated reactions, raising the possibility that extracellular GSH directly counteracts copper-catalyzed cysteine oxidation. We first identified a concentration of CuCl_2_ sufficient to oxidize cysteine within 1 hour (Fig. S3), then incubated 100nM or 200nM CuCl_2_ with or without 1mM GSH (Fig. 2e). GSH suppressed copper-driven cysteine oxidation at both copper concentrations, accompanied by consumption of GSH and accumulation of cysteinyl-glutathione (Cys-GSH) and oxidized glutathione (GSSG) that increased with copper (Fig. 2e). Because copper catalyzes glutathione oxidation and thiol– disulfide interconversion, these changes reflect a redox interaction between GSH and copper and do not, on their own, distinguish reduction of cystine from copper chelation. However, the quantity of GSH required far exceeded what chelation would demand. Keeping cysteine fully reduced in the presence of dFBS required ∼1 mM GSH (Fig. 2f), roughly 2,000-fold more than the total copper present (∼500 nM, largely protein-bound). Following the reaction over time, GSH reduced cystine to cysteine through a Cys-GSH intermediate (Fig. 2g), which was itself resolved by further exchange with GSH to release cysteine (Fig. 2h). These data indicate that GSH acts primarily as a cystine reductant through disulfide exchange (Fig. 2i). Thus, at the 1.1mM concentration measured in the lung TME, GSH can both protect existing cysteine and drive the cystine-to-cysteine equilibrium via disulfide exchange, making cysteine the predominant bioavailable species in vivo.

### Tissue context and stromal cells establish the reduced microenvironment

The chemistry above establishes that sustaining extracellular cysteine requires a continuous supply of reducing equivalents, raising the question of where in the tumor those originate. This reduced state could reflect an intrinsic property of the cancer cells or a feature imposed by the surrounding tissue. To distinguish these possibilities, we injected lung tumor cells (Kras^G12D/+^; Trp53^fl/fl^) subcutaneously into C57BL/6 NJ mice, placing genetically identical cells in a non-native tissue context (Fig. 3a). In sharp contrast to the autochthonous lung TME, the interstitial fluid of the resulting subcutaneous tumors was dominated by cystine (19μM) and oxidized glutathione, and total glutathione (GSH + GSSG) was significantly lower than in GEMM tumors (Fig. 3b). Thus, the tissue in which the tumor arises is a critical determinant of TME redox composition, pointing to a stromal or tissue-resident source of reducing equivalents.

**Figure 3:**
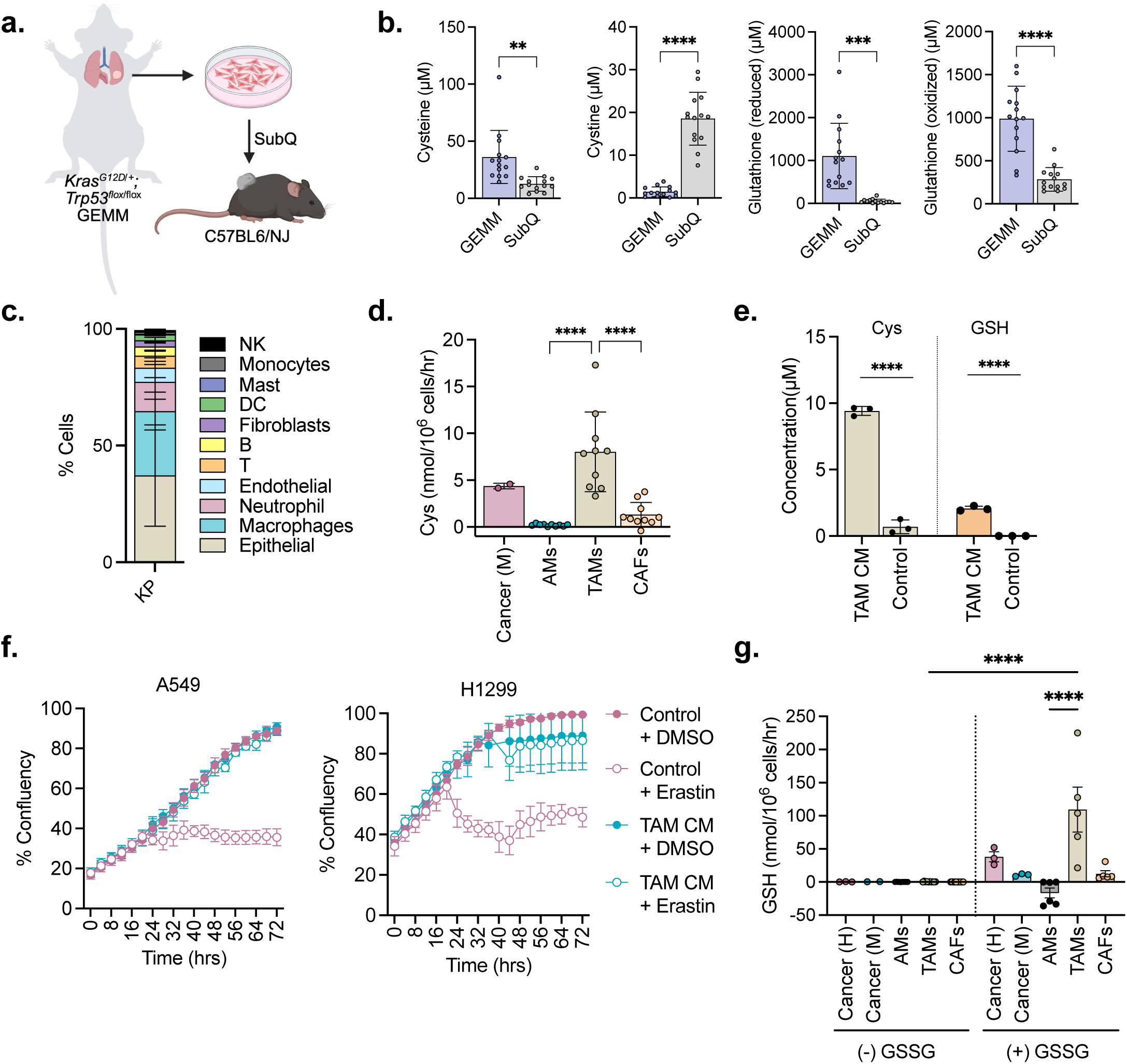
Tumor associated macrophages actively create and maintain the reduced microenvironment. **(a)** Schematic of subcutaneous tumor generation in C57BL/6 NJ mice, using cells isolated and cultured from a LSL-Kras^G12D^; Trp53^flox/flox^ lung tumor with a C57BL/6 J background. **(b)** Concentration of cysteine, cystine, and reduced and oxidized glutathione in interstitial fluid of subcutaneous and GEMM tumor. n = 14 for subcutaneous and GEMM mice groups, GEMM data are from Figure 1 for comparison. **(c)** Frequency of the indicated cell populations in lung tumors from mice harboring LSL-Kras^G12D^; Trp53^flox/flox^ mutations (n = 3), determined by single cell analysis. **(d)** Rate of cysteine release from murine lung cancer cells (n = 2), alveolar macrophages from healthy mice (AMs, n = 10), tumor alveolar macrophages (TAMs, n = 10) and cancer-associated fibroblasts (CAFs, n = 10) from tumor-bearing mice. **(e)** Quantification of cysteine and GSH in TAM-conditioned medium and matched cell-free control medium used in **(f).** n = 3 independent replicates. **(f)** Growth of H1299 and A549 cells incubated in TAM-conditioned medium or matched control medium, with DMSO or 5 µM (H1299) or 10 µM (A549) Erastin. Cell growth was monitored by live-cell imaging (CellCyteX). n = 5 independent replicates. Data are representative of 2 independent experiments. **(g)** Rate of glutathione accumulation in the conditioned media from human cancer cells (n = 3), mouse cancer cells (n = 1), alveolar macrophages from healthy mice (AMs, n = 6), tumor alveolar macrophages (TAMs, n = 5) and cancer-associated fibroblasts (CAFs, n = 5) from tumor-bearing mice in the presence and absence of 910 µM GSSG. All data are mean ± s.d. Statistical analysis was performed by Welch’s test (**b**), One-way ANOVA with Tukey’s multiple comparison test (**d,g**), or unpaired t test (**e**).

To identify that source, we analyzed single-cell RNA sequencing data from Kras^G12D/+^; Trp53^fl/fl^ lung tumors, which showed that after epithelial cells (37.2%), macrophages (27.6%) and neutrophils (12.5%) are the major non-malignant cell populations in the lung TME, with fibroblasts being a minor population (2.7%) (Fig. 3c). To test the contribution these populations directly, we profiled conditioned medium from murine cancer cells, alveolar macrophages from healthy mice (AM), alveolar macrophages from tumor-bearing mice (TAM), and cancer-associated fibroblasts (CAFs) by LC-MS. Neutrophils did not survive isolation and short-term culture, precluding analysis of their conditioned medium. Given the rapid oxidation of free cysteine in serum-containing media (Fig. 2a), conditioned media were collected in the presence of TCEP, allowing extracellular cysteine to be quantified directly. TAMs released cysteine at substantially higher rates than cancer cells, AMs or CAFs (Fig. 3d), with TAM-conditioned media containing 9.4 µM cysteine (Fig. 3e).

We next asked whether TAM-derived cysteine is sufficient, on its own, to support lung cancer cell growth when cystine import is inhibited. Cysteine titration in A549, H1299, and H2009 cells showed that 10 µM cysteine could sustain growth and delay cell death following xCT inhibition with erastin, with confluence increasing and death decreasing in a dose-dependent manner (Fig. S4). Consistent with this, TAMs-conditioned medium, but not matched cell-free control medium, sustained proliferation of A549 and H1299 cells during erastin treatment (Fig. 3f). Basal GSH levels were only in ∼2 µM TAM conditioned media (Fig. 3e), and secretion rates were low across all cell populations (Fig. 3g), but adding GSSG at the concentration measured in the lung TME drove substantially greater conversion to extracellular GSH by TAMs than cancer cells, AMs or CAFs (Fig. 3g). Together, these findings show that lung tumor-associated macrophages both secrete cysteine and reduce GSSG to GSH, thereby promoting a reduced extracellular environment.

### A physiological sulfur medium alters cellular metabolic dependencies

Cysteine-supplemented RPMI does not recapitulate the broader nutrient context of the TME, including its distinct glutamate, GSH and taurine content, which can influence cellular cysteine handling via effects on xCT activity and cysteine demand, respectively. To more directly model the *in vivo* redox microenvironment, we compared the sulfur metabolite and amino acid composition of lung tumor IF with standard RPMI (Fig. 4a) and formulated a sulfur tumor interstitial fluid medium (sTIF) matching the measured physiological concentrations. sTIF supported normal cell growth and fully prevented erastin-induced death in H1299, H2009 and A549 cells over 72 hours (Fig. 4b, Fig. S5a). Cells grown in sTIF had substantially lower intracellular cysteine than cells in standard RPMI (5-fold lower in A549 and H2009 and 2-fold lower in H1299), yet intracellular GSH was maintained (H2009) or even slightly elevated (A549, H1299) (Fig. 4c). Intracellular cystine was also higher in RPMI than in either cysteine- or sTIF-cultured cells (Fig. S5b). Together, these results show that the intracellular cysteine pool tracks extracellular cyst(e)ine availability, which is supraphysiological in standard RPMI. The physiological sulfur context of sTIF supports both growth and glutathione maintenance despite a much smaller free cysteine pool.

**Figure 4:**
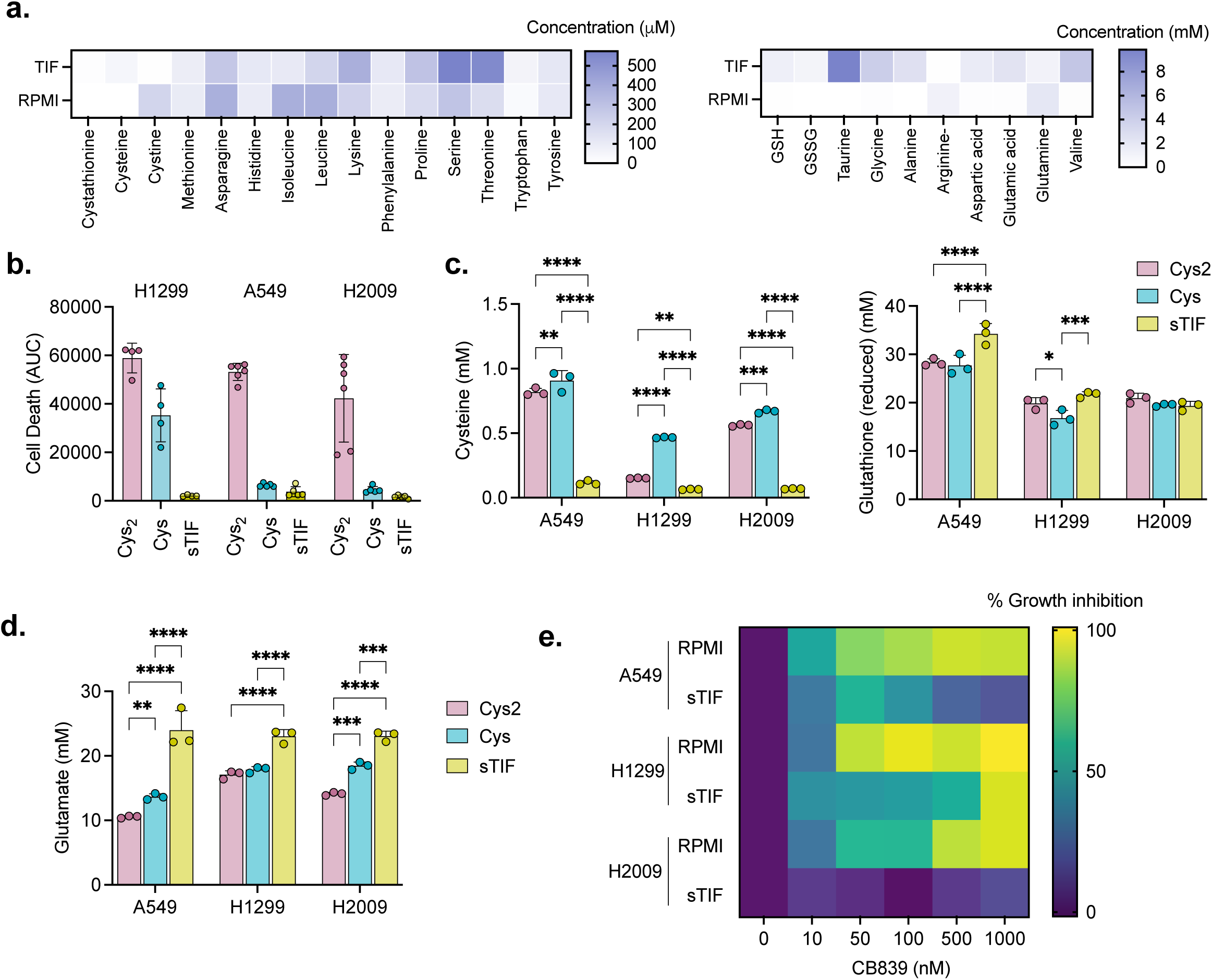
Physiological culture conditions confer cystine independence. **(a)** Concentrations of the indicated metabolites in RPMI vs. lung tumor interstitial fluid (TIF). **(b)** Quantification of cell death in H1299, H2009, and A549 cells cultured in cysteine-containing medium, cystine-containing medium, or lung sTIF medium. Cell death was quantified as the area under the curve (AUC) of SYTOX Green-positive cells normalized to cell confluence over 72 h of live cell imaging. **(c,d)** Intracellular cysteine and glutathione (**c**) and glutamate (**d**) concentration in A549, H1299 and H2009 cells cultured in RPMI under cysteine (Cys: 200µM cysteine and 100µM TCEP), cystine (Cys_2_: 200µM cystine), and sTIF conditions for 8 h. All conditions contain 10% dialyzed FBS (dFBS). **(e)** Heatmap showing percent inhibition of cell growth in presence of the indicated concentration of CB-839 in A549, H1299 and H2009 cells cultured in RPMI or sTIF media + 10% dFBS for 72 h. Data are mean ± s.d. n = 4 (**b**), n = 3 (**c,d**), or n = 5 (**e**) biological replicates, with at least 2 (**c-e**) or 3 (**b**) independent experiments. Statistical analysis was done by two-way ANOVA followed by Tukey’s multiple comparison test.

Because reduced reliance on cystine import predicts reduced need for glutamate that xCT exports in exchange, we asked whether cells grown in sTIF accumulate glutamate and become less dependent on glutaminolysis. Indeed, we found cells in cysteine conditions accumulate glutamate, which was even more pronounced in sTIF media (Fig. 4d). Correspondingly, cells in sTIF were more resistant to the glutaminase inhibitor CB839, remaining proliferative at doses up to 1µM CB839 treatment in A549 and H2009 and 0.5µM in H1299 (Fig. 4e, Fig S5c). Together, these data show that the physiological sulfur amino acid and metabolite composition of the TME, rather than cysteine alone, impacts cellular metabolism and alters metabolic dependencies.

### Physiological conditions promote resistance to oxidative stress

Next, we asked whether the reduced TME alters how cells respond to oxidative stress. Measuring both basal and menadione-inducible reactive oxygen species (ROS), we found that basal ROS did not differ significantly between RPMI and sTIF, whereas menadione-induced ROS accumulation was blunted in sTIF across cell lines (Fig. 5a), suggesting sTIF enhances the cells’ capacity to buffer an oxidative insult. To test whether this buffering translated into protection from cell death, we challenged cells across RPMI and sTIF with a panel of oxidants and ferroptosis inducers acting through distinct mechanisms and assessed viability. sTIF conferred resistance to the superoxide-generating compound β-lapachone, and the lipid peroxidation inducer cumene hydroperoxide, evident as reduced cell death relative to RPMI (Fig. 5b,c, Fig. S6a,b). sTIF similarly conferred resistance to the GPX4 inhibitors RSL3 and ML162 in both human and murine lung cancer cells lines (Fig. 5d,e, Fig. S6c,d). To avoid any potential confounding interaction of our drugs with sTIF components, we generated H1299 cells expressing D-amino acid oxidase (D-AAO), which oxidizes D-alanine to produce H_2_O_2_ inside the cell. These cells died when cultured in RPMI media with 5-10mM D-alanine, but not the equivalent L-alanine, and this death was prevented when cells were instead cultured in sTIF with the same D-alanine concentration (Fig. 5f, Fig. S6e). Because GSH levels were similar between media conditions (Fig. 4c), we asked whether oxidative stress protection instead reflects a difference in NADPH availability. While the steady state NADPH/NADP+ ratio was not altered by sTIF medium (Fig. 5g), sTIF medium preserved NADPH levels during menadione or ML162 exposure, which declined progressively in RPMI-cultured H1299 and H2009 cells (Fig. 5h,i). Together, these results indicate that cancer cells are more resistant to oxidative stress in sTIF than in standard culture.

**Figure 5:**
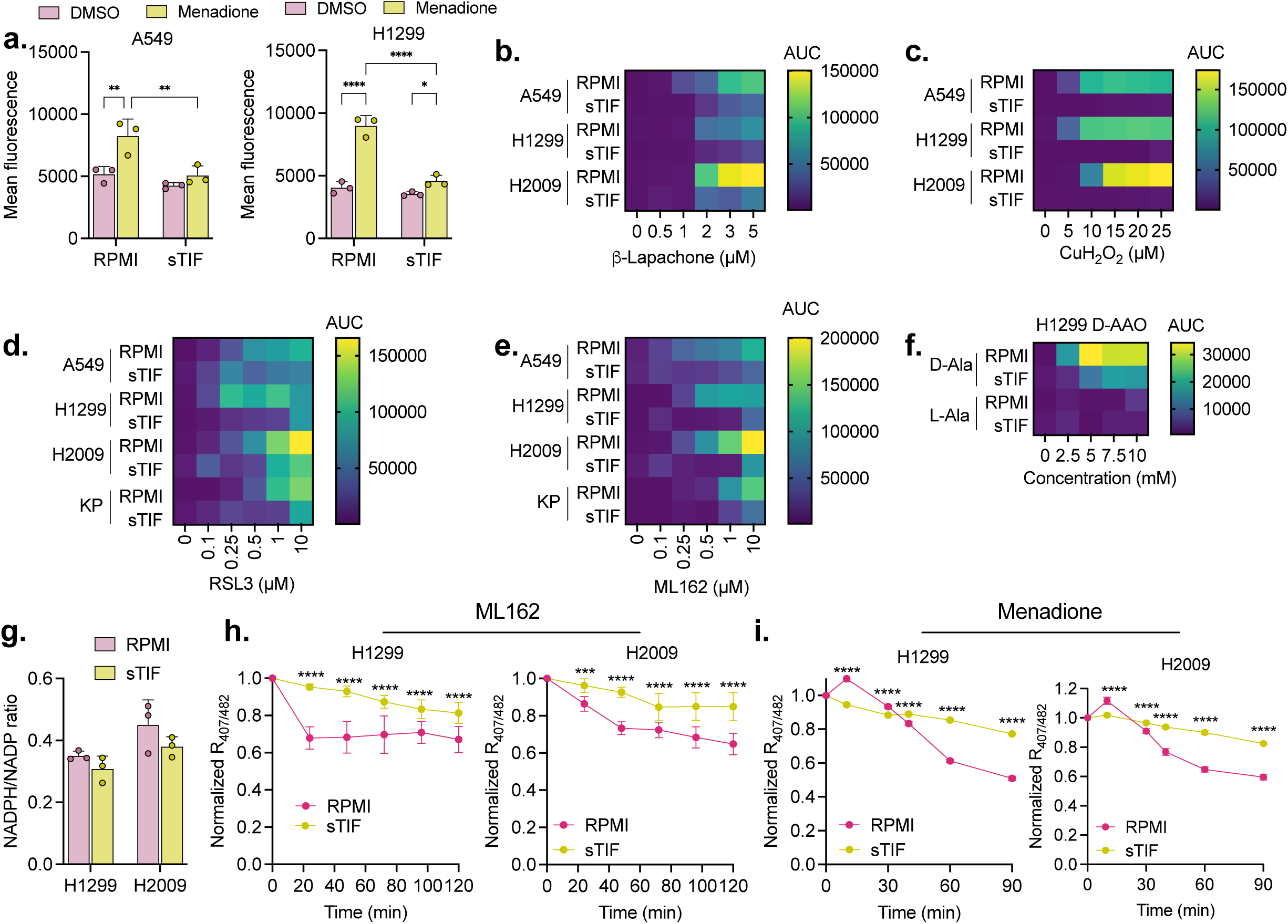
Physiological conditions promote resistance to ROS. **(a)** Intracellular ROS in A549 and H1299 cells cultured in RPMI or sTIF, measured with CellROX after 30 minutes exposure to DMSO or 10µM menadione. All media contained 10% dialyzed FBS (dFBS). **(b-f)** Heatmap showing relative cell death at the indicated concentrations of β-lapachone (**b**), cumene hydroperoxide (CuH_2_O_2_) (**c**), RSL3 (**d**) or ML162 (**e**) in A549, H1299, H2009 and murine KP cells, and of D-Alanine and L-Alanine in D-Amino acid oxidase (D-AAO)-expressing H1299 cells (**f**), cultured in RPMI or sTIF medium + 10% dFBS over 64-72 h. **(g)** Intracellular NADPH/NADP ratio in H1299 and H2009 cells cultured in RPMI or sTIF medium for 2 h. All conditions contain 10% dialyzed FBS (dFBS). **(h-i)** NADPH availability indicated by normalized ratio (407nm/482nm) in iNap1-Cyto expressing H1299 and H2009 cells in response to 1µM ML162 (**h**) or 10µM menadione (**i**) cultured in RPMI or sTIF media for the indicated time points. For **a, g, h, i**, data are mean ± s.d. n = 3 (**a,g**) n = 10 (**h, i**), or n = 5 (**b-f**) biological replicates, with at least 2 independent experiments. Statistical analysis was done by two-way ANOVA followed by Fisher’s Least Significant Difference (LSD) test (**a**), or two-way ANOVA with Sidak’s multiple comparison test (**h-i**).

### Extracellular cysteine renders GSR and TXNRD1 dispensable

If the TME supplies cysteine already in reduced form, cells should require less intracellular reductase capacity to generate it. We therefore asked whether cysteine availability alters the requirement for GSR and TXNRD1. We generated GSR-knockout (GSR KO) A549 and H1299 cells and confirmed loss of GSR protein expression by western blot (Fig. 6a). In standard RPMI, GSR KO cells were more sensitive than control cells to the TXNRD1 inhibitor auranofin (Fig. 6b; Fig. S7a). This sensitivity was reversed by culturing cells in lung sTIF, which shifted the auranofin dose-response curve of GSR KO cells to the right (Fig. 6b; Fig. S7a). A comparable rescue was observed when cells were cultured in cysteine-containing medium stabilized with TCEP (Fig. 6b; Fig. S7b). These results show that extracellular reduced cysteine, whether supplied in sTIF media or stabilized via TCEP, diminishes cancer cell reliance on intracellular cystine reduction mediated by GSR and TXNRD1.

**Figure 6:**
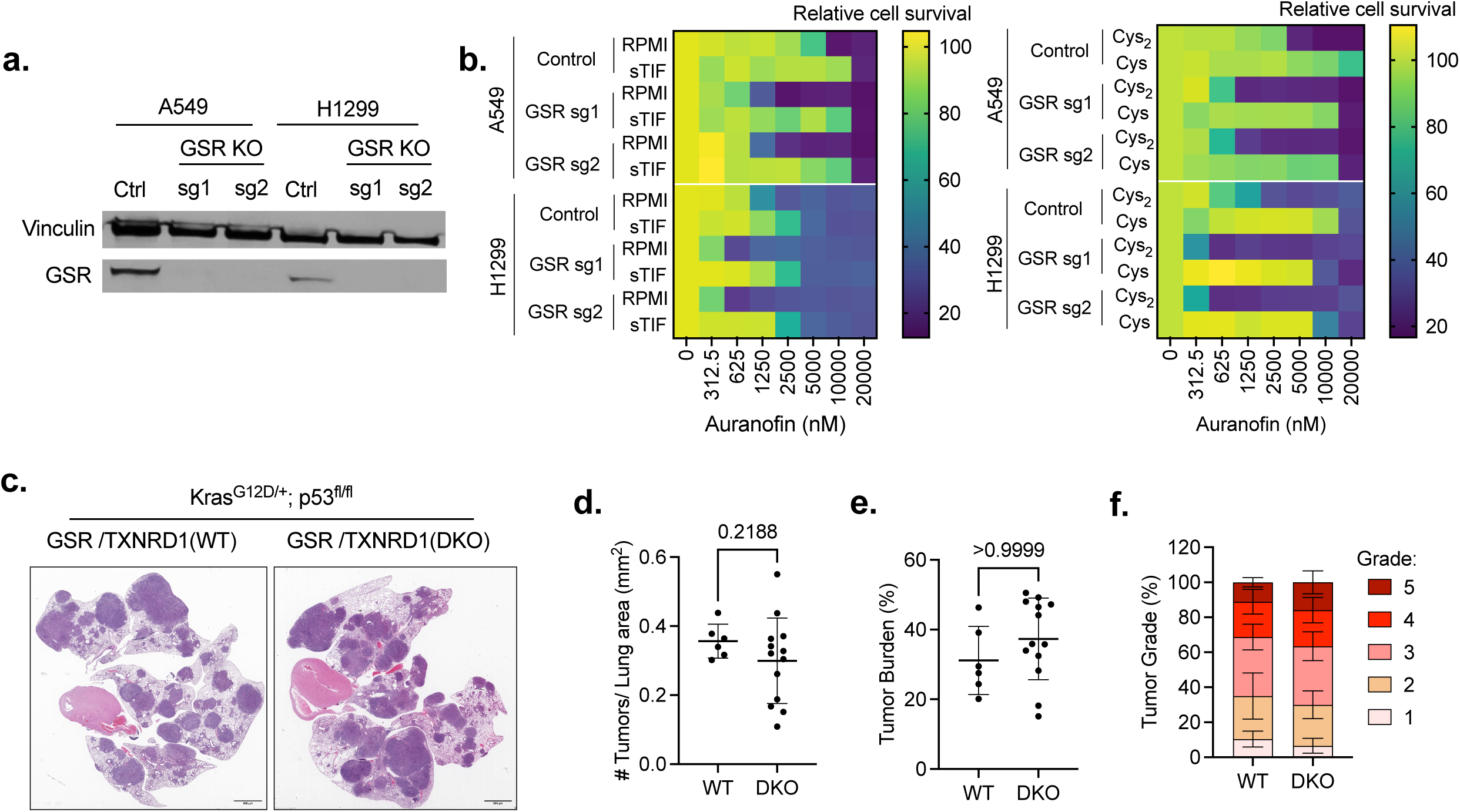
Cysteine uptake confers independence from cytosolic disulfide reductases. **(a)** Immunoblot of GSR expression in control and GSR-knockout A549 and H1299 cells. Vinculin was used as a loading control. **(b)** Survival of control and GSR-knockout A549 and H1299 cells cultured in cysteine (Cys: 200µM cysteine and 100µM TCEP), cystine (Cys_2_: 200µM cystine), and sTIF conditions and treated with the indicated concentrations of auranofin. All conditions contain 10% dialyzed FBS (dFBS). Survival is shown as percent of confluence relative to the no auranofin condition at each concentration. Data are representative of at least 3 independent experiments. **(c)** Representative hematoxylin and eosin-stained lung sections from Kras^G12D/+;^ p53^fl/fl^ mice bearing GSR/TXNRD1 wild-type or GSR/TXNRD1 double-knockout tumors. Scale bars 500µm. **(d)** GSR/TXNRD1 wild-type (n=6) and double-knockout (n=13) tumor number normalized to total lung area. **(e)** GSR/TXNRD1 wild-type (n=6) and double-knockout (n=13) tumor burden (total tumor area/total lung area). **(f)** Distribution of GSR/TXNRD1 wild-type (n=6) and double-knockout (n=13) histological tumor grades. For **d-f**, data are mean ± s.d. Statistical analysis was done by Wilcoxon test.

To test whether this relationship holds in vivo, we compared lung tumors from *Kras^G12D/+^; Trp53^fl/fl^* mice that were wild-type or double knockout for *Gsr* and *Txnrd1*. Double knockout tumors developed readily and were histologically indistinguishable from wild-type tumors (Fig. 6c), with no significant difference in tumor number (Fig. 6d), tumor burden (Fig. 6e), or tumor grade distributions between genotypes (Fig. 6f). Together, these results demonstrate that the sulfur metabolite composition of the tumor microenvironment renders lung cancer cells independent from cytosolic glutathione and thioredoxin reductases.

## Discussion

Previous work has characterized the nutrient composition of interstitial fluid in tumors and tissues^10,11^. Here, we focus specifically on the metabolites that maintain redox homeostasis, and our collection and extraction methods preserve their native redox state. The nutrient composition of tumor interstitial fluid differed markedly from that of either the healthy organ of origin or serum, most notably in its sulfur metabolites. This difference implies that nutrients available to transformed cells during initiation, whether supplied from the originating niche or the circulation, differ from those available as the tumor develops, and that both cancer cell–intrinsic factors and the various cell populations of the tumor microenvironment (TME) actively shape that environment. Our most striking observation was that cysteine was more abundant than its oxidized counterpart, cystine, in the TME, a finding that challenges the prevailing model in which cells acquire cysteine by importing cystine and reducing it intracellularly.

Reconciling our observations with the well-established role of xCT requires consideration of the cell types in which the transporter operates. Non-small cell lung cancer expresses high levels of xCT, which is correlated with advanced stage^13^, and NRF2 activity^14^. This establishes that the transporter is expressed and active, but not that cystine uptake is the primary source of cellular cysteine. Even direct tracing is ambiguous on this point. We previously traced cystine and observed labeling of tumor cysteine pools^9^, which we took as evidence of cystine import through xCT but because cystine and cysteine interconvert by disulfide exchange in the circulation, that labeling is equally consistent with cysteine import. Pharmacological evidence also carries the same ambiguity. xCT Inhibition with sulfasalazine and HG106 slow tumor growth in the KP model, but also inhibit xCT in non-malignant cells, including macrophages, and may act by starving tumors of stroma-derived cysteine rather than by blocking cystine import into cancer cells^15^. While acute Slc7a11 deletion in established PDAC induces ferroptosis and extends survival, xCT was also deleted in the stromal and immune compartments^8^. Deletion confined to the tumor compartment had no effect and targeting xCT in both PDAC cells and CAFs was required to see an effect in orthotopic models^16^. Wherever non-malignant cells import cystine and release cysteine, targeting xCT will deplete the extracellular pool to impact the reduced microenvironment. Because it depletes both species systemically, engineered cyst(e)inase^7^ offers a therapeutic strategy that does not rely on which one cancer cells import or on the compartment that supplies it.

The reduced state of the lung TME raises the question of how it is maintained. We find that when cancer cells grow in a non-native tissue environment, the interstitial fluid of the resulting tumor is composed of oxidized species, indicating that the reduced state is not cell-intrinsic but is instead contributed by the tissue in which the tumor arises. We find that tumor-associated macrophages secrete cysteine at higher rates than either cancer cells or alveolar macrophages from healthy lung, pointing to tumor-specific remodeling. Macrophage cysteine release in response to lipopolysaccharide or TNF has been reported previously^17^. Macrophages can also export cysteine through a cystine- and glucose-dependent process involving cell-surface thioredoxin reductase, at least under pharmacological induction^18^. Glutathione secretion by pulmonary alveolar macrophages is also increased following exposure to particulates^19^. We find macrophages further contribute to the extracellular redox environment by reducing extracellular GSSG to GSH, although whether this reflects surface-localized reductase activity, or uptake and intracellular reduction followed by export, remains unresolved. The model that emerges is one in which cancer cells remodel resident macrophages, which in turn sustain the redox environment of the TME by secreting cysteine and reducing available GSSG.

To test the consequences of this environment, we formulated sTIF, a medium containing sulfur metabolites and amino acids at the concentrations measured in LUAD interstitial fluid. Its combined cysteine and cystine content is 44 µM (95% cysteine), which is sufficient to support proliferation, and because cells take up both species with equal efficiency, they become cystine independent. The glutamate demand that xCT imposes contributes to glutamine addiction^5,20,21^, and glutaminase inhibition has been pursued therapeutically but has not succeeded beyond preclinical studies^22,23^. Cystine independence and the high intracellular glutamate in the tumor microenvironment can both contribute to glutaminase inhibitor resistance and offer an explanation for those failures. sTIF also conferred resistance to oxidants and ferroptosis inducers. Although baseline GSH levels and NADPH/NADP^+^ ratios were comparable between RPMI and sTIF, cells in sTIF maintained their NADPH levels under oxidative challenge. The reduced demand on the thioredoxin and glutathione reductase systems that accompanies cystine independence may leave greater NADPH capacity to counter intracellular ROS, and the concomitant glutamate accumulation may further support NADPH regeneration by feeding the malate and citrate pools that supply malic enzyme (ME1) and IDH1. These observations have implications for oxidative stress as a therapeutic strategy^24^, since radiation and many chemotherapeutics act through ROS and a reduced TME would blunt their effect. More broadly, the shifts in metabolic dependency between physiologically matched and standard media suggest that a redox-matched medium such as sTIF could be used for drug or CRISPR screening under more physiological conditions.

Several questions remain open. Interstitial fluid measurements report a bulk average across the tumor, yet a microenvironment maintained by macrophage-derived reducing equivalents is unlikely to be spatially uniform. Regions distant from macrophage infiltrate may remain oxidized and lead to heterogeneity in xCT dependence and responses to xCT inhibition. Testing this will require spatially resolved measurement of thiol redox state, which the centrifugation-based methods used here cannot provide. The transporter responsible for cysteine uptake in LUAD also remains undefined. Although our tracing data show that cells take up cysteine and cystine with comparable efficiency on a per-cysteine-equivalent basis, we have not established which of the transporters reported to carry cysteine is responsible. Direct cysteine import has been studied most thoroughly in the brain, where neuronal uptake proceeds largely through EAAT3 (SLC1A1)^25^, but the corresponding route in peripheral tissues and tumors is unclear. SLC1A4 (ASCT1) and SLC1A5 (ASCT2) have both been proposed^26,27^, but their role in cysteine uptake is yet to be studied, and recent work indicates that cysteine acts as a modulator of SLC1A5 rather than as a substrate^28^. Finally, cells cultured in sTIF maintained intracellular cysteine two- to five-fold lower than cells in standard RPMI while sustaining or increasing GSH, indicating that conventional culture substantially overestimates the size of the intracellular free cysteine pool. Because cysteine availability governs coenzyme A synthesis, iron-sulfur cluster assembly, hydrogen sulfide production^1^, and cysteine’s own toxicity to cultured cells^29–31^, the physiological set point for each of these processes may differ from what culture-based measurements suggest.

Our study has several limitations. We examined three adenocarcinoma models, so whether the reduced microenvironment is a general feature of solid tumors or a property specific to epithelial tumors arising in specific tissue contexts remains unresolved. Mesenchymal, neuroendocrine, and hematological malignancies may behave differently, especially given how widely stromal composition varies across these settings. All interstitial fluid measurements were made in mouse models, and the associated concentrations in patient tumors are unknown. Interstitial fluid collection is also low-yield and required pooling material from multiple tumors, so the concentrations we report are averaged across lesions that necessarily obscure both intratumoral spatial organization and inter-tumor heterogeneity.

## Methods

### Generation of experimental animals

All animal experiments were approved by the University of South Florida Institutional Animal Care and Use Committee (protocols 6358R, 7922R, 8736R and 12540R). Mice were housed under 12:12 hour (light/dark cycle) conditions at 22°C (+/-2°C), group-housed, with access to standard rodent chow diet, at the Moffitt Stabile Research Building animal facility. Sample sizes for each experiment are indicated in the corresponding figure legends.

#### Hepatocellular carcinoma (HCC) model

Mouse models were generated as previously described^9^. Briefly, Myc; p53^-/-^ HCC was induced in 8-week-old C57BL/6 NJ female mice (Jackson laboratory, RRID: IMSR_JAX:005304) by hydrodynamic tail vein injection. Each mouse received 10µg of pT3-EF1A-MYC-IRES-luc plasmid (addgene number: 129775), 10µg of pX330 p53 plasmid (addgene number: 59910) and 2.5µg of CMV-SB13 plasmid^32^, diluted in sterile saline (Hospira, Catalog # NDC 0409-4888-02) to a volume equal to 10% of body weight. Injections were performed via the lateral tail vein over 7 seconds using a 27-gauge needle, with mice restrained without anesthesia. Tumors developed approximately 4–6 weeks post-injection and were monitored by bioluminescence imaging. Mice received 150 mg/kg D-luciferin via i.p. injection and were imaged on a IVIS Lumine S5 imaging system (Revvity, CLS148588) at 3 weeks post injection to confirm successful DNA delivery.

#### Pancreatic ductal adenocarcinoma (PDAC) model

LSL-Kras^G12D/+^; Trp53^flox/flox^; p48-cre mice were generated by intercrossing LSL-Kras^G12D^ (RRID: IMSR_JAX:008179), Trp53^flox^ (RRID: IMSR_JAX:008462) and p48-cre (RRID: IMSR_JAX:023329) lines. Mice developed tumors with a median survival of 60 days and tumor development was monitored by abdominal palpation.

#### Lung adenocarcinoma (LUAD) model

LSL-Kras^G12D/+^; Trp53^flox/flox^ mice were generated by intercrossing LSL-Kras^G12D^ and Trp53^flox^ lines (RRIDs as above). At 4-8 weeks of age, tumor formation was initiated by intranasal installation of 2x10^7^ plaque-forming unit (PFU) Ad5CMVCre (University of Iowa) in a total volume of 50 μL under 2% isoflurane anesthesia. Mice developed tumors with a median survival of 3-4 months and were used for experimentation when they showed evidence of tumor burden, including rapid respiration.

#### Subcutaneous tumor model

A cell line derived from a lung tumor of a female LSL-Kras^G12D^; Trp53^flox/flox^ mouse (C57BL/6J background) was resuspended in a 1:1 PBS:Matrigel solution (1x10^6^ cells/100μL) and injected into the flank subcutaneous region of 9-10 weeks old female C57BL/6 NJ mice. Tumor growth was monitored by calipers every alternative day, with volume calculated as (length × width× height)/2. Once tumors reached ∼300mm^3^ (approximately 4 weeks post-injection), tumors were collected.

### Single-cell RNA-sequencing analysis

Single-cell RNA sequencing, data processing, filtering, batch effect correction, clustering, and cluster annotation were performed as described previously^33^, with the following modification: following cluster annotation, the four dendritic cell subtypes (cDC1, cDC2, mregDC, and pDC) were aggregated into a single dendritic cell (DC) population for downstream analyses.

### QuPath analysis and tumor grading

H&E-stained slides were prepared by IDEXX Bioanalytics and scanned at 20x magnification using an Aperio AT2 digital slide scanner (Leica Biosystems). Whole-lung sections were analyzed in QuPath by an investigator blinded to genotype. The total lung area and individual tumor lesions were manually annotated, and each tumor was assigned a histopathological grade from 1 to 5 according to previously established grading criteria for the *Kras*^G12D^ lung tumor model^34^. Tumor number density was calculated by dividing the total number of tumors in each H&E section by the corresponding total lung area. Tumor burden was calculated by dividing the combined area of all annotated tumors by the total lung area. To determine the distribution of tumor burden across histological grades, the combined area of tumors assigned to each grade was divided by the total tumor area for that mouse and expressed as a percentage.

### Serum collection

Blood (100-200µl) was collected into BD Microtainer SST Blood Collection Tubes (Cat #365967) from the cheek vein prior to sacrifice and centrifuged at 10,000 rpm for 15 minutes at 25°C. The supernatant (serum) was collected, aliquoted into cryovials, and snap-frozen in liquid nitrogen. Serum was stored at -80°C until analysis.

### Interstitial fluid collection

Mice were sacrificed via cervical dislocation, and tumor or organ samples were briefly rinsed with room-temperature sterile saline (Fisher Scientific, Catalog # Z1376) to remove residual blood, then blotted dry with Whatman or Kimwipes. Tissue was placed on a 20μm nylon mesh filter (Repligen, Catalog # 888-13801-PK) affixed to a 50ml conical tube, capped, and centrifuged at 310 x g for 10 minutes at 4°C. The interstitial fluid that collected at the bottom of the tube was immediately snap-frozen in liquid nitrogen. For pooled organ samples, pancreases from three mice were combined per sample during collection, and interstitial fluid from liver or lung from three mice were combined post collection. Samples were stored at -80°C prior to metabolomics processing.

### Metabolomics sample preparation

#### Liquid samples (serum, TIF, IF, and media)

Samples were diluted in extraction buffer (80% methanol containing 0.5 mmol/L ammonium formate, 5 mmol/L N-ethyl maleimide [NEM], and internal standards, pH 7) to the following final dilutions: 1:10 for serum, 1:20 for media, and 1:10-1:100 for TIF/IF depending on sample volume. Internal standards were added at the concentrations listed below:

| Isotope-labeled internal standard | Formula | Source/Catalog No. | Extraction buffer concentration (µM) |
| --- | --- | --- | --- |
| Cysteine Sulfinic acid-<sup>13</sup>C<sub>3</sub><sup>15</sup>N | <sup>13</sup>C<sub>3</sub>H<sub>7</sub><sup>15</sup>NO<sub>4</sub>S | Prepared in-house | 0.5 |
| Cysteine-S-sulfate-d3 | C<sub>3</sub>H<sub>4</sub>D<sub>3</sub>NO<sub>5</sub>S<sub>2</sub> | CDN isotopes D-6386 | 1 |
| L-Cystathionine-d4 | C<sub>7</sub>H<sub>10</sub>D<sub>4</sub>N<sub>2</sub>O<sub>4</sub>S | Cambridge Isotope Laboratories, DLM-6108 | 1 |
| Cystine-d4 | C<sub>6</sub>H<sub>8</sub>D<sub>4</sub>N<sub>2</sub>O<sub>4</sub>S<sub>2</sub> | Cambridge Isotope Laboratories, DLM-9812 | 1 |
| Glutathione disulfide-<sup>13</sup>C<sub>4</sub><sup>15</sup>N<sub>2</sub> | C<sub>16</sub><sup>13</sup>C<sub>4</sub>H<sub>32</sub>N<sub>4</sub><sup>15</sup>N<sub>2</sub>O<sub>12</sub>S<sub>2</sub> | Cambridge Isotope Laboratories, CNLM-8782 | 1 |
| Hypotaurine-d4 | $C_2H_3D_4NO_2S$ | CDN isotopes<br>D-5158 | 1 |
| Methionine-$^{13}C_5$ | $^{13}C_5H_{11}NO_2S$ | Cambridge Isotope<br>Laboratories,<br>CLM-893-H | 1 |
| N-Acetyl-Methionine-d3 | $C_7H_{10}D_3NO_3S$ | CDN isotopes<br>D-7678 | 1 |
| Ophthalmic acid-$^{13}C_2, ^{15}N$ | $C_9^{13}C_2H_{19}N_2^{15}NO_6$ | Millipore sigma<br>739308 | 1 |
| S-(5'-Adenosyl)-L-Homocysteine-d4 | $C_{14}H_{16}D_4N_6O_5S$ | CDN isotopes<br>D-6452 | 1 |
| S-(5'-Adenosyl)-L-Methionine-d3 | $C_{15}H_{20}D_3N_6O_5S$ | CDN isotopes<br>D-4093 | 1 |
| Taurine-$^{13}C_2$ | $^{13}C_2H_7NO_3S$ | Cambridge Isotope<br>Laboratories,<br>CLM-6622 | 1 |
| Cysteinylglycine-NEM-d5 | $C_{11}H_{12}D_5N_3O_5S$ | Prepared in-house | 1 |
| Cysteamine-NEM-d5 | $C_8H_9D_5N_2O_2S$ | Prepared in-house | 0.1 |
| L-Cysteine-NEM-d5 | $C_9H_9D_5N_2O_4S$ | Prepared in-house | 0.1 |
| Gamma-glutamylcysteine-NEM-d5 | $C_{14}H_{16}D_5N_3O_7S$ | Prepared in-house | 1 |
| Glutathione-NEM-d5 | $C_{16}H_{19}D_5N_4O_8S$ | Prepared in-house | 1 |
| L-Homocysteine-NEM-d5 | $C_{10}H_{11}D_5N_2O_4S$ | Prepared in-house | 0.1 |
| N-Acetyl-L-Cysteine-NEM-d5 | $C_{11}H_{11}D_5N_2O_5S$ | Prepared in-house | 0.1 |
| Pantetheine-NEM-d5 | $C_{17}H_{24}D_5N_3O_6S$ | Prepared in-house | 1 |
| Coenzyme A-NEM-d5 | $C_{27}H_{38}D_5N_8O_{18}P_3S$ | Prepared in-house | 1 |

After addition of extraction buffer, samples were vortexed and incubated for 30 min at 4°C, then centrifuged at 13,000 x g for 10 minutes at 4°C. The supernatant was collected and stored at - 80°C until LC-MS analysis.

#### Cell samples

Cells were seeded in 6-well plates in at least biological triplicate at a density of 0.2 million/well and extracted at the indicated time points. Media was aspirated and cells were washed with DPBS on wet ice. Ice-cold extraction buffer (400µl; 80% methanol containing 0.5 mmol/L ammonium formate, 5 mmol/L NEM, and internal standards, pH 7) was then added, and plates were incubated for 30 min at 4°C. Cells were scraped, and the supernatant was collected after centrifugation (13,000 x g, 4°C, 10 minutes) and stored at -80°C until LC-MS analysis.

### Internal standard preparation for thiol-containing metabolites

Thiol (-SH)-containing standards – cysteamine, homocysteine, N-acetyl-cysteine, L-cysteine, glutathione, cysteinylglycine, gamma-glutamylcysteine, pantetheine and Coenzyme A – were each prepared as 10mM stock solutions in 10% Formic acid. 5mM N-ethylmaleimide-d5 (NEM-d5; CDN isotopes, D-6141) was prepared in 10mM ammonium formate (pH 7). For each standard, 100µl of the 10mM stock was added to 900µl of 5mM NEM-d5 solution and incubated at room temperature for 30min to yield a 1mM derivatized internal standard.

### LC/MS analysis and data processing

Metabolite quantification was performed by LC-MS technology as previously described^30^. In brief, chromatographic separation was performed on a Vanquish UPLC system (Thermo Fisher Scientific) equipped with a Waters Atlantis Premier BEH z-HILIC VanGuard FIT column (2.1mm ID x 150mm, 2.5µm particle size; Waters), held at 30°C. A 5 µl sample volume was injected via autosampler. Mobile phase A (10 mmol/L ammonium carbonate and 0.05% ammonium hydroxide in water) and mobile phase B (100% acetonitrile) were used with the following gradient: 0–13 minutes, 80% to 20% of B; 13–15 minutes, 20% of B; 15–20 minutes, 80% of B. Metabolites were detected using a Q Exactive HF (QE-HF) Orbitrap mass spectrometer (Thermo Fisher Scientific) with heated electrospray ionization (H-ESI), scanning m/z 65–950 in both positive and negative modes. The following instrument conditions were maintained: capillary temperature: 320°C, voltage: 3.5 kV, mass resolution: 120,000, and AGC target: 3x10^6^. Raw data (.raw files) were analyzed in Skyline^35^ (version 23.1.0.268). Metabolites were identified by matching exact precursor ion m/z (within a 10ppm window) and retention time against an in-house library of authentic standards. Peak area was used as a read-out of metabolite intensity, normalized to internal standards and corrected for dilution factor to determine concentration. For isotope correction, the extracted metabolite signals from Skyline were loaded into the IsoCor^36^ (Version 2.2.3) and for dual-isotope correction, a custom correction matrix incorporating natural isotope abundances of ^13^C (1.07%) and ^34^S (4.29%) was applied in Excel.

### Cell growth and death assay

Cells were seeded in 96-well plates in RPMI + 10% dFBS, with 3-5 replicates per condition, at the following seeding densities: A549, 3,000 cells/well; H1299, 8-10,000 cells/well; H2009, 5,000 cells/well; and murine KP, 3,000 cell/well. After overnight incubation at 37°C, media was aspirated, cells were washed with cystine-, methionine- and glutamine-free RPMI (US Biological Life Sciences, R9016) and then grown in the specified experimental media formulations, supplemented with 20nM SYTOX Green (Invitrogen) at 37°C in CellCyteX live cell imager (Cytena). Images (4 per well) were acquired every 4 hours for 72-96 hours and processed using CELLCYTE Studio to quantify confluency (proliferation) and SYTOX Green-positive events (cell death). Cell death was expressed as the SYTOX Green-positive signal normalized to confluency. For some cell death experiments, the area under the curve (AUC) was calculated for each replicate as the sum of the normalized SYTOX Green/confluency values across all imaged time points. For analysis of percent growth inhibition, the following formula was used: [(AUC of Control - AUC of Experimental)/AUC of Control] *100.

To assess the effect of TAM-conditioned medium on erastin-induced cell death, A549 and H1299 cells were seeded as described above with the following modifications: cells were seeded at 2,500 (A549) and 10,000 (H1299) cells/well in 100 µl complete RPMI 1640 medium and following the cystine/methionine/glutamine-free media wash, treated with 100 µl of TAM-conditioned medium or matched cell-free control medium in the presence or absence erastin (5 µM for H1299 and 10 µM for and A549). The treatment medium was replaced every 24 hours for 72 hours and imaging was performed on the CellCyteX.

### NADPH and NADPH/NADP ratio analyses

The NADPH/NADP ratio was measured using the NADP/NADPH-Glo Assay kit (Promega, G9082). Cells were seeded in 6-well plates with 3 replicates per condition, at 0.2x10^6^ cells/well in RPMI + 10% FBS. After overnight incubation at 37°C, media was aspirated, cells were washed with cystine-, methionine- and glutamine-free RPMI (US Biological Life Sciences, R9016), and cells were cultured in the specific experimental media formulations at 37°C for 2 hours before processing according to manufacturer protocol and luminescence analysis on a GloMax Discover plate reader (Promega).

NADPH availability was assessed using a genetically encoded fluorescent biosensor, iNap1-Cyto, which was optimized to report on the cytosolic NADPH pool^37^, stably expressed in cells following lentiviral transduction. For each experiment, 150,000 cells were seeded overnight on a 6cm dish and then pretreated with either RPMI control or sTIF media conditions for 2 hours prior to being challenged with the indicated treatment. Biosensor fluorescence was captured with an Axiovert 5 (Zeiss) fluorescent microscope fitted with Colibri 3 LED light source to permit excitation of iNap1-Cyto at 407nm and 482nm. Multichannel images were analyzed with ZEN microscopy software (Zeiss) to determine the fluorescence intensity of individual cells at each excitation wavelength and then the ratio (407nm/482nm) determined to indicate NADPH availability. For each experiment, fluorescence images of the same field of view were captured prior to and following the media pretreatment and then for the duration of the treatment challenge; the fluorescence ratio of at least 10 distinct cells was assessed for each condition.

### Intracellular ROS analysis

The intracellular ROS levels were measured using CellROX Green (Invitrogen, Waltham, MA, USA). Cells were seeded at 1.5x10^5^ cells/well in 24-well plates in RPMI + 10% FBS, with three replicate wells per condition, and incubated overnight incubation at 37°C. The media was then aspirated, and cells were washed with cystine-, methionine- and glutamine-free RPMI (US Biological Life Sciences, R9016) before incubation in either RPMI or sTIF for 1 hour at 37°C. Then media was then replaced with RPMI or sTIF containing 5µM CellROX Green for 30 minutes, after which menadione was added to a final concentration of 10µM for a further 30 minutes. Cells were harvested by trypsinization and analyzed on a BD Accuri C6 Plus flow cytometer using the FITC channel. Data were analyzed with FlowJo software (BD Biosciences). Mean fluorescence intensity (MFI) was calculated for each sample, and the MFI of unstained cells was subtracted as background.

### Statistical analysis

Statistical analyses were performed using GraphPad Prism (version 10.1.2). P < 0.05 was considered statistically significant (*, P < 0.05, **, P < 0.01, ***, P < 0.001, ****, P < 0.0001). Welch’s t-test was used to compare two groups. One-way ANOVA with Tukey’s multiple comparisons test was used to compare a single variable across three or more groups. Two-way ANOVA with Tukey’s multiple comparisons test was used to compare two variables. Data are presented as mean ± SD, and the number of independent animals or primary cell lines (n) or experimental replicates for each analysis is indicated in the figure legends.

### Cell Culture

Human lung adenocarcinoma cell lines H1299, A549, and H2009 were obtained from the Hamon Cancer Center Collection at the University of Texas Southwestern Medical Center or commercially from the American Type Culture Collection, as previously described^38^. Cells were maintained in RPMI 1640 medium (Gibco, 11875-093) supplemented with 10% fetal bovine serum (FBS; Sigma-Aldrich, F0926), without antibiotics. Murine KP cells were also maintained in RPMI 1640 supplemented with 10% GBS without antibiotics. Lenti X 293T cells (Clontech, 632180) were cultured in DMEM (Fisher Scientific, MT10013CV) supplemented with 10% FBS. Macrophages were maintained in RPMI 1640 supplemented with 10% heat-inactivated FBS, 1 mM sodium pyruvate (Corning, 25-000-CL), 20 ng/ml recombinant mouse GM-CSF (Miltenyi Biotec, 130-095-746), and 1% penicillin–streptomycin (Gibco, 15070-063). CAFs were maintained in DMEM medium supplemented with 10% FBS. All cultures were incubated at 37 °C in a humidified atmosphere of 95% air and 5% CO₂. Mycoplasma contamination was routinely monitored using the MycoAlert Mycoplasma Detection Kit (Lonza, LT07-418).

For experiments involving manipulation of cysteine and cystine availability, cells were cultured in RPMI 1640 medium lacking L-glutamine, L-methionine, L-cysteine, and L-cystine (US Biological, R9016), prepared according to the manufacturer’s instructions. This deficient medium was supplemented with 10% dialyzed FBS (Sigma-Aldrich, F0392), 100 µM L-methionine (Sigma-Aldrich, M5308), 2 mM L-glutamine (Gibco, 25030-081), and L-cysteine (Sigma-Aldrich, 168149) and/or L-cystine (Sigma-Aldrich, C6727), as specified for each experiment. Additional reagents utilized in these experiments included dimethyl sulfoxide (DMSO; VWR, 97063), erastin (Selleckchem, S7242), reduced glutathione (GSH; Sigma-Aldrich, G4251), tris(2-carboxyethyl)phosphine hydrochloride (TCEP; Sigma-Aldrich, C4706), bathocuproine disulfonic acid disodium salt hydrate (BCS; Sigma-Aldrich, 146625), and auranofin (MedChemExpress, HY-B1123).

### Preparation of lung tumor interstitial fluid like medium

Lung sulfur tumor interstitial fluid-like medium (sTIF) was prepared using L-glutamine-, L-methionine-, L-cysteine-, and L-cystine-free RPMI 1640 powder (US Biological, R9016) prepared according to the manufacturer’s instructions. The medium was sterilized by filtration through a 0.22µm membrane. Individual nutrient stocks were prepared in ultrapure water unless otherwise indicated. L-cysteine (Sigma-Aldrich, 168149), reduced glutathione (GSH; Sigma-Aldrich, G4251), and oxidized glutathione (GSSG; Sigma-Aldrich, G4376) were dissolved in 1 N HCl. L-cystine (Sigma-Aldrich, C6727) was dissolved in 2 N HCl, and L-aspartic acid (Sigma-Aldrich, 194633) was dissolved in 0.5 M HCl. L-cystathionine (Sigma-Aldrich, C7505), L-methionine (Sigma-Aldrich, M5308), taurine (Sigma-Aldrich, T0625), glycine (Fisher Scientific, G46-1), L-glutamic acid (Sigma-Aldrich, G8415), and L-valine (Sigma-Aldrich, V0513) were dissolved in ultrapure water. Stock solutions were aliquoted and stored at −80 °C to avoid repeated freeze-thaw cycles. Immediately before use, the base medium was supplemented as in this table:

| Metabolite | Concentration ($\mu\text{M}$) | Metabolite | Concentration (mM) |
| --- | --- | --- | --- |
| Cystathionine | 6 | Glutathione | 1.36 |
| Cysteine | 42 | Taurine | 10 |
| Cystine | 2 | Glycine | 4 |
| Glutathione disulfide | 910 | Aspartic acid | 1.6 |
| Methionine | 90 | Glutamic acid | 2.3 |
|  |  | Glutamine | 2 |
|  |  | Valine | 4.7 |

Following supplementation, the pH was adjusted to 7.4. Dialyzed FBS was then added to a final concentration of 10%, and the completed medium was sterile filtered through 0.22µm filter. Lung sTIF medium was prepared freshly on the day of each experiment and was not stored.

### Lentiviral transduction

Lentivirus was produced by transfecting Lenti X 293T cells with 6 μg of the lentiviral plasmid and 6 μg packaging plasmids pCMV-VSV-G (Addgene, 8454) or pMD2.G (VSV-G, Addgene, 12259) and pCMVdR8.2 dvpr (Addgene, 8455) or psPAX2 (Addgene, 12260) at a 1:8 ratio, using PEI (polyethylenimine) transfection reagent (Sigma-Aldrich, 408727). The following plasmids were used: D-AAO [pLGH-DAAO-NES-Hyg, cloned from pC1-CMV-DAAO-NES (Addgene #141107)]; iNap1-Cyto [pLenti-CMV-Hygro-iNap1-Cyto, cloned from a construct purchased from FR Biotechnology] both were cloned into the pLenti-CMV-Hygro backbone (Addgene, 17446). The sgRNAs were cloned into plentiCRISPR v2 for the following target sequences: control: GAAAAGACTATTTCAAGCAGA; GSR sg1: TCAGTGATGTCTTAGAACCC; GSR sg2: GGGCTTGGGATCACTCGTGA. Viral supernatant was collected at 24-hour intervals and diluted 1:1 with complete medium, supplemented with polybrene at a final concentration of 8μg/mL (Santa Cruz Biotechnology, sc-134220A), and used to transduce target cells overnight. Transduced cells were selected with 250μg/mL Hygomycin (Invivogen, ant-hg-1) or 2 μg/mL puromycin. D-AAO expression was confirmed by cell death in 10mM and 20mM D-Alanine (Thomas Scientific LLC, C916D39) conditions but not L-Alanine (Acros Organics, A0407465). GSR knockout was confirmed by Western blot.

### Isolation of Murine Lung Cancer Cells

Lung cancer cells were isolated from Kras^G12D^; p53^flox/flox^ lung tumor bearing mice on a pure C57BL/6J background. Tumors were macrodissected from the lung and digested in 2mg/ml Collagenase/Dispase (Roche, 10269638001) in Advanced DMEM/F12 (Gibco, 11320033), cells were pelleted and washed in complete DMEM, and cells were plated in matrigel (Corning, CB-40230) and overlaid with Advanced DMEM/F12 + 10% FBS + 1x ITS (Gibco, 41400045) + 0.5µM A83-01 (Tocris, 2939) + 10.5µM Y-27632 (Cayman, 10005583) to establish organoid cultures. After organoids were passaged several times, organoids were adapted to 2D conditions, followed by adaptation to RPMI + 10% FBS.

### Isolation of Alveolar Macrophages

Alveolar macrophages were isolated from 4-month-old C57BL6 NJ mice (AMs) and Kras^G12D^; p53^flox/flox^ lung tumor bearing mice (TAMs) by bronchoalveolar lavage as described previously^39^. In brief, mice were euthanized by cervical dislocation. A 20-gauge cannula was inserted into the trachea towards the lungs. The lungs were lavaged with 1 ml of pre-warmed bronchoalveolar lavage (BAL) buffer [PBS (VWRL0119), 2 mM EDTA (Invitrogen, AM9260), 0.5% heat-inactivated fetal bovine serum (FBS)]. The BAL buffer was slowly instilled and withdrawn. This was repeated 10 times with fresh BAL buffer, and the recovered fluid was pooled. The pooled BAL fluid was filtered through a 70µm cell strainer and cells were collected by centrifugation at 300xg for 5 min. Residual red blood cells were lysed by ACK lysis buffer (Gibco, A10492-01). The cells were washed in BAL buffer, and viable cells were seeded in 5 ml of complete macrophage medium supplemented with 20 ng/ml recombinant mouse GM-CSF at a maximum of 400K cells into one well of a six-well plate. The culture medium was replaced 12-18 hours after plating to remove nonadherent cells, and the remaining macrophages were maintained for subsequent experiments.

### Isolation of Cancer-associated Fibroblasts

Mouse lung tumors were freshly collected, minced using a sterile razor blade, and digested in 20 ml DMEM containing 1 mg/ml collagenase type IV (Gibco, 17104-019), 0.1% soybean trypsin inhibitor (Gibco, 17075-029), 50 U/ml DNase I (Fisher Scientific, NC0421464), and 0.125 mg/ml dispase (Gibco, 17105-041) for 30 minutes at 37 °C with agitation, mixing every 10 minutes. The digested tissue was centrifuged at 300xg for 3 minutes, and the resulting pellets were resuspended in 10 ml prewarmed trypsin-EDTA (Corning, 25-052-CI) and incubated for 10 minutes at 37 °C with agitation. Trypsinization was stopped by adding 10 mL cold DMEM + 10% FBS, and the cell suspensions were filtered through 100 µm cell strainers. Cells were collected by centrifugation, washed once with DMEM, and incubated in 1 ml ACK lysis buffer for 3-5 minutes at room temperature. Red blood cells lysis was stopped by adding 9 mL DMEM, followed by centrifugation to collect the cells.

Cancer associated fibroblasts (CAFs) were sorted by magnetic-activated cell sorting or fluorescence-based cell sorting. For magnetic-activated cell sorting, cells were resuspended in 100 µl complete DMEM and incubated with 2 µl biotin-conjugated anti-PDGFRα antibody (BioLegend, 135910) for 15 minutes. Cells were washed with PBS and then incubated with 10 µl Streptavidin MicroBeads (Miltenyi Biotec, 130-048-101) in 90 µl separation buffer (PBS supplemented with 0.5% bovine serum albumin and 2 mM EDTA) for 10 minutes at 4 °C protected from light. Following an additional wash, cells were resuspended in 500 µl separation buffer and applied to MS columns positioned in an OctoMACS separator. Columns were washed with 500 µl separation buffer, then removed from the magnetic separator, and the retained PDGFRα-positive cells (CAFs) were eluted with 500 µl separation buffer. For fluorescence-based sorting, cells were resuspended in phenol red-free DMEM supplemented with 10% FBS. Fc receptors were blocked with anti-mouse CD16/CD32 antibody (BD Biosciences, 553142; 1:20) for 15 minutes at room temperature. Cells were stained with biotin-conjugated anti-PDGFRα antibody (BioLegend, 135910; 0.25 µg/ million cells) and FITC-conjugated anti-CD31 antibody (BioLegend, 102405; 1:150) for 30 min in the dark. After washing, cells were incubated with APC-conjugated streptavidin (BioLegend, 405207; 1:500) for 30 minutes. DAPI was added at 100 ng/ml to exclude nonviable cells. Unstained and single-color controls were included for compensation and gate determination. DAPI-negative, CD31-negative, PDGFRα-positive cells were isolated using a MACSQuant Tyto cell sorter with a standard cartridge and collected into complete phenol red-free DMEM. CAFs isolated using either OctoMACS Separator (Miltenyi Biotec) or a MACSQuant Tyto Cell Sorter (Miltenyi Biotec) were seeded in DMEM + 10% FBS at a maximum of 500K cells into one well of a six-well plate.

### Conditioned medium collection

Conditioned medium was collected to quantify extracellular cysteine and glutathione. Cells were seeded in 24-well plates in 1 mL complete medium per well and grown until 70-80% confluent. Then, the medium was removed and replaced with 1 ml fresh medium containing 10% dFBS + 100 µM TCEP. Matched cell-free wells containing the same volume of medium were included as controls for each condition. For experiments testing the effect of oxidized glutathione, medium was prepared with or without 910µM GSSG. Plates were incubated for 8 h (LC-MS) or 24 h (erastin protection) at 37 °C and 5% CO₂. After collection, conditioned media was immediately placed on dry ice and stored at -80 °C until use. For LC-MS experiments, cells were detached following medium collection and counted for normalization. Metabolite levels in matched cell-free medium controls were subtracted from conditioned media samples and normalized to the final cell number.

### Immunoblotting

Cells cultured in 10cm plates were washed once with ice-cold PBS and lysed in 500 µl RIPA buffer (Sigma-Aldrich, SIAL-20-188) supplemented with 0.1% (w/v) SDS (Sigma-Aldrich, 74255) and Pierce Protease Inhibitor Mini Tablets (Thermo Fisher Scientific, PI88666). Cells were scraped into the lysis buffer, incubated on ice for 30 min, and centrifuged at maximum speed for 10 min to remove insoluble material. Protein concentrations were determined using the DC Protein Assay (Bio-Rad, 5000113–5000115) according to the manufacturer’s instructions.

20µg protein from each sample was prepared in 6× SDS sample buffer and separated on NuPAGE 4–12% Bis-Tris gels (Invitrogen, NP0321BOX) in MOPS SDS running buffer (Invitrogen, NP0001). Proteins were transferred to 0.45-µm Amersham Protran nitrocellulose membranes (Cytiva, 10600003). The membrane was blocked in 5% non-fat dry milk (CellPro, MB9696) prepared in Tris-buffered saline containing 0.1% Tween 20 (TBST) for 1 h at room temperature and incubated overnight at 4 °C with antibodies against GSR (Santa Cruz Biotechnology, sc-133245, 1:1000) and vinculin (Sigma-Aldrich, V9264, 1:10000). Goat anti-mouse IgG secondary antibody (JacksonImmunoResearch, 115-035-003) was used. Protein bands were visualized using Clarity ECL substrate (Bio-Rad, 1705060) or a luminol-based homemade ECL substrate.

### Inductively coupled plasma mass spectrometry (ICP-MS)

Copper concentrations in RPMI medium with or without 10% dialyzed fetal bovine serum (dFBS) were measured by ICP-MS. Medium samples (10 µl) were diluted in 290 µl of 4% trace-metal-grade nitric acid (TraceSELECT; Sigma-Aldrich, 84385) prepared in ultrapure water. Samples were incubated for 2 hours at room temperature and centrifuged at 15,000 rpm for 10 min to pellet insoluble material. The clarified supernatants were collected for analysis. Samples were analyzed using an iCAP TQ ICP-MS instrument (Thermo Fisher Scientific). Copper was quantified using the ^63^Cu isotope. A 4% nitric acid reagent blank was included, and calibration standards containing 10, 100, 500, and 1,000 ppb copper were prepared from a 10-ppm multi-element standard containing copper (ICP Multi-Element Standard Mix 5, CMS-5). Copper concentrations were determined from the calibration curve and adjusted for the 30-fold dilution introduced during sample preparation.

## Acknowledgements

We would like to thank Mara Sherman for advice about CAF isolation and Ryan Martin for assistance with ICP-MS. This work was supported by grants from the NIH/NCI (R01CA309659 and P01CA250984) to G.M.D. This work was also supported by the Moffitt Flow Cytometry, Microscopy and Proteomics/Metabolomics Cores, which are funded in part by Moffitt’s Cancer Center Support Grant (NCI, P30CA076292).

## Contributions

G.M.D., F.K.C. and B.A.Y. conceived the study. F.K.C. and B.A.Y. designed experiments and interpreted the results with input from G.M.D. F.K.C. performed metabolomic profiling of interstitial fluid. F.K.C. and B.A.Y performed metabolomics analyses of cultured cells and media. B.A.Y. developed the sTIF medium, isolated tumor-associated macrophages and CAFs, performed primary culture experiments, and analyzed tumor burden. F.K.C. performed the drug sensitivity assays and oxidative stress and ferroptosis experiments. N.P.W. performed NADPH assays with F.K.C. Y.H. and C.J. generated GSR knockout cell lines and the D-AAO construct. S.A-A. and E.R.F. performed scRNA-seq. M.L. and M.C. performed animal experiments. P.G-S., M.L. and P.N.S. collected interstitial fluid. G.M.D. supervised the study and acquired funding. F.K.C., B.A.Y. and G.M.D. wrote the manuscript with input from all authors.

## Supplementary Tables and Figures

**Supplementary Table 1: Concentrations of metabolites in interstitial fluid and serum.** Concentrations of metabolites in tumor interstitial fluid from tumors (tab 1), healthy tissue (tab 2), and serum concentrations from tumor models (tab 3).

**Supplementary Figure 1:**
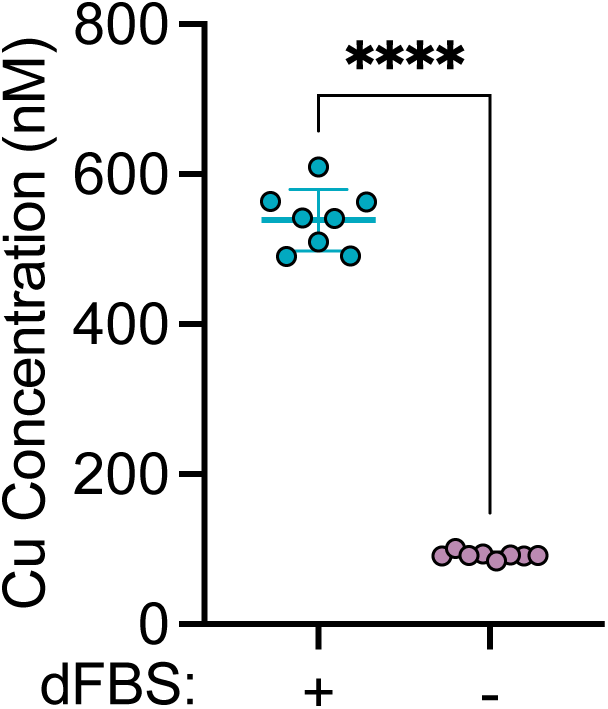
Impact of serum on copper concentration in RPMI media. Copper concentration in RPMI medium with or without 10% dialyzed FBS (dFBS), measured by ICP-MS. Data are mean ± s.d.; Statistical analysis was done by Welch’s test. n = 8 biological replicates. Data are representative of 2 independent experiments.

**Supplementary Figure 2:**
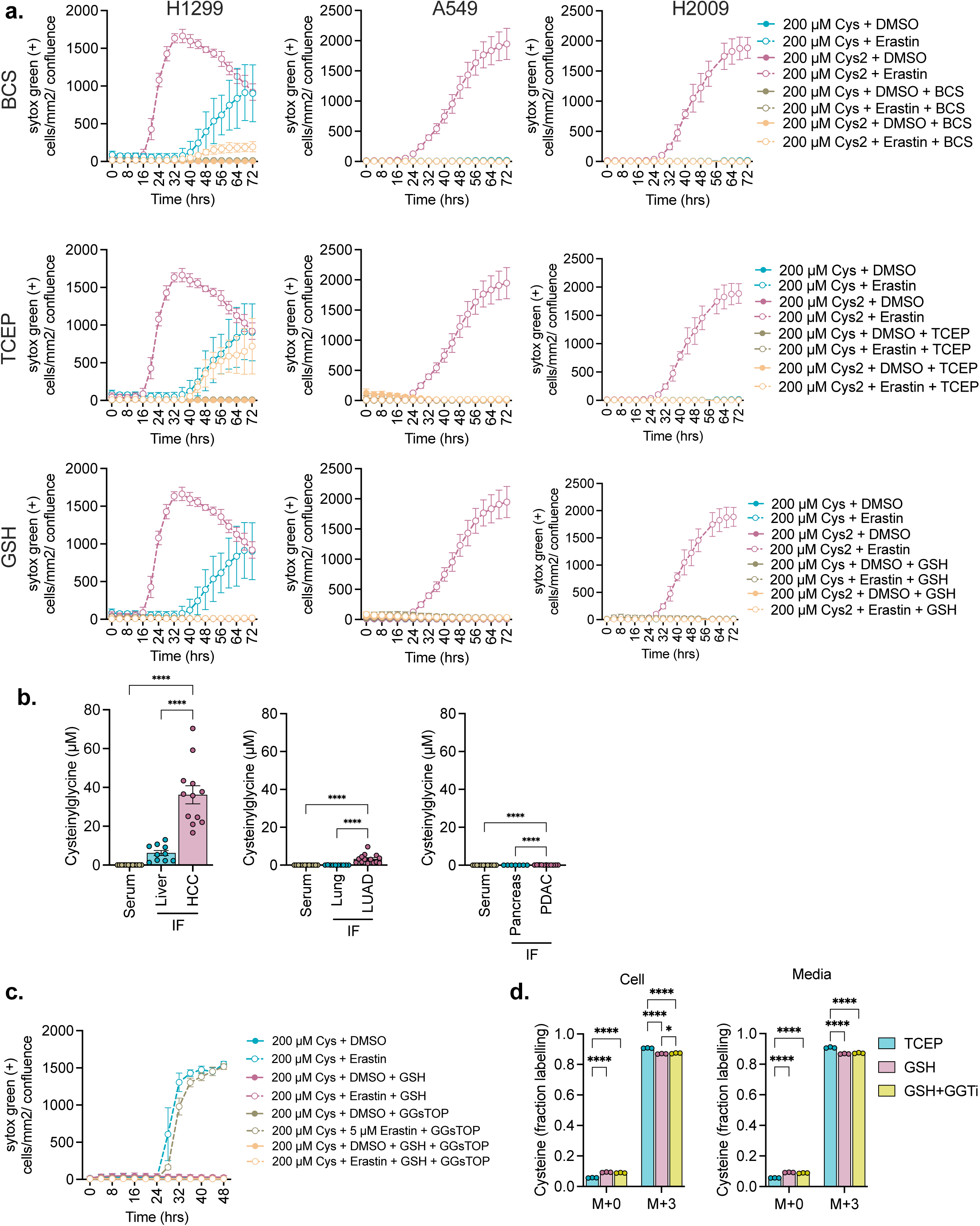
Cysteine stabilizing conditions alter Erastin-induced cell death in lung cancer cells. **(a)** H1299, A549, and H2009 cells were cultured in cysteine or cystine-containing medium (200 µM) supplemented with 20 µM BCS, 100 µM TCEP, or 1mM GSH, with or without Erastin (5 µM for H1299; 10 µM for A549 and H2009). SYTOX Green-positive cells normalized to confluence (cell death) over time (72 h). **(b)** Concentration of cysteinylglycine in interstitial fluid of the indicated healthy organ and tumor, and serum from matched tumor-bearing mice. Data are mean ± s.d.; Liver: n = 12 (HCC), 10 (liver), 11 (serum). Lung: n = 14 (LUAD), 10 (lung), 14 (serum). Pancreas: n = 11 (PDAC), 7 (pancreas), 11 (serum). **(c)** H1299 cells were cultured in cysteine-containing medium (200 µM) supplemented with or without 1mM GSH, with or without 5 µM Erastin. SYTOX Green-positive cells normalized to confluence (cell death) over time (48 h). **(d)** H1299 cells were cultured in [U-^13^C]-cysteine-containing medium (200 µM) supplemented with 100µM TCEP or 1mM GSH or 1mM GSH with 1µM GGTi for 24 hours with media being changed after 12 hours. For **a,c,** and **d** data are mean ± s.d.; n = 5 (**a, c**) or n = 3 (**d**) biological replicates. Data are representative of 3 (**a**), 2 (**c**) or 1 (**d**) independent experiment. Statistical analysis was performed by one-way ANOVA with Tukey’s multiple comparisons test (**b,d**).

**Supplementary Figure 3:**
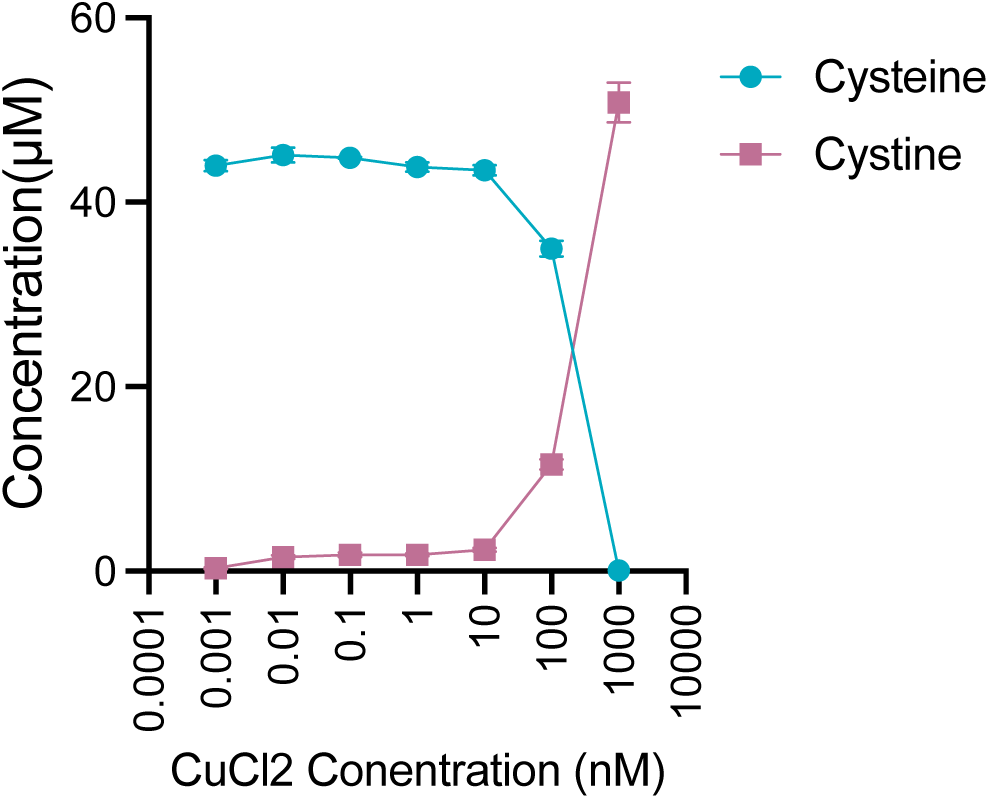
CuCl_2_ mediated cysteine oxidation. Cysteine and cystine levels following incubation of 100µM cysteine with the indicated CuCl_2_ concentrations in serum-free RPMI media for 1 hour. Data are mean ± s.d.; n = 3 biological replicates, and 1 independent experiment.

**Supplementary Figure 4:**
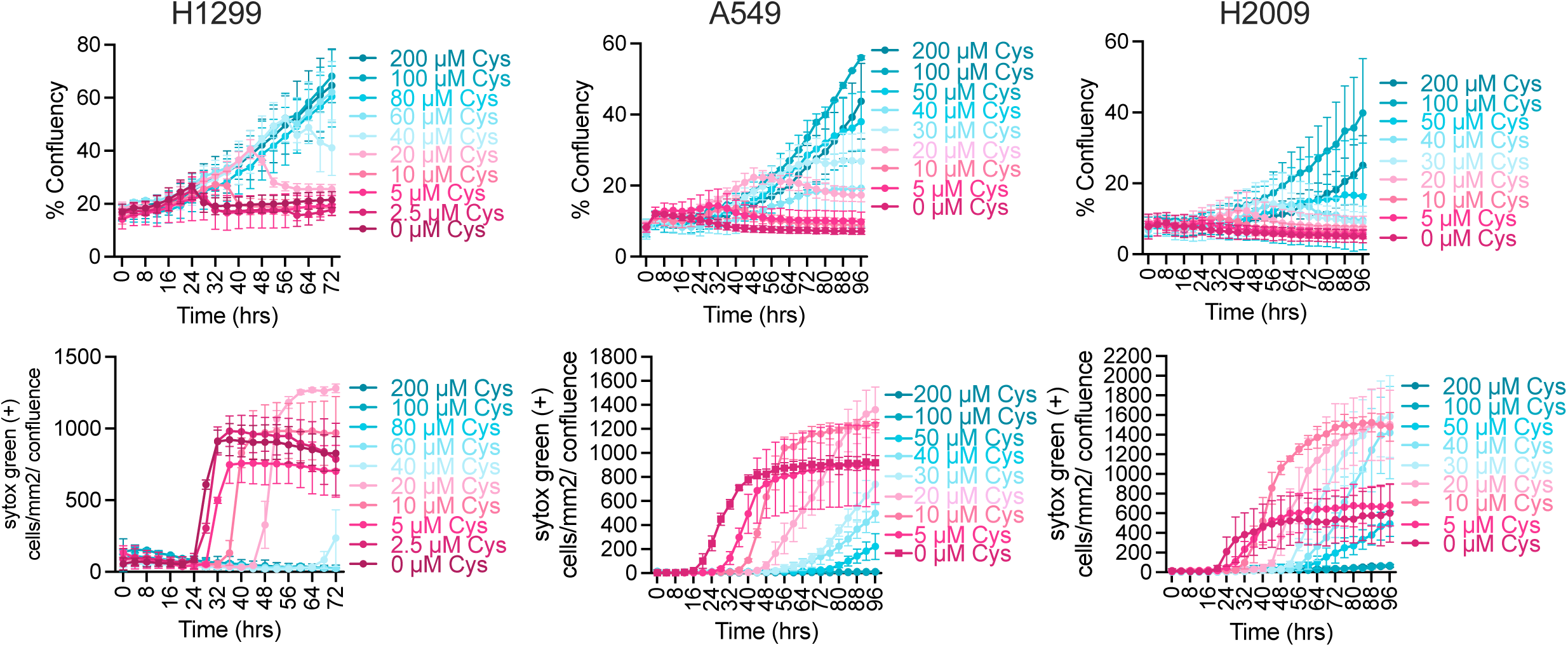
Cysteine supports lung cancer cell growth and suppresses cell death in a dose-dependent manner. H1299, A549, and H2009 cells were cultured in RPMI + dFBS supplemented with Erastin (5 µM for H1299; 10 µM for A549 and H2009) and the indicated concentrations of cysteine stabilized with 100 µM TCEP. Cell growth and death were monitored by live-cell imaging (CellCyteX). (top) Percent confluence over time. (bottom) SYTOX Green-positive cell counts, normalized to confluence, over time. Data are mean ± s.d.; n = 3 biological replicates. Data are representative of 2 (H2009) or 3 (A549, H1299) independent experiments.

**Supplementary Figure 5:**
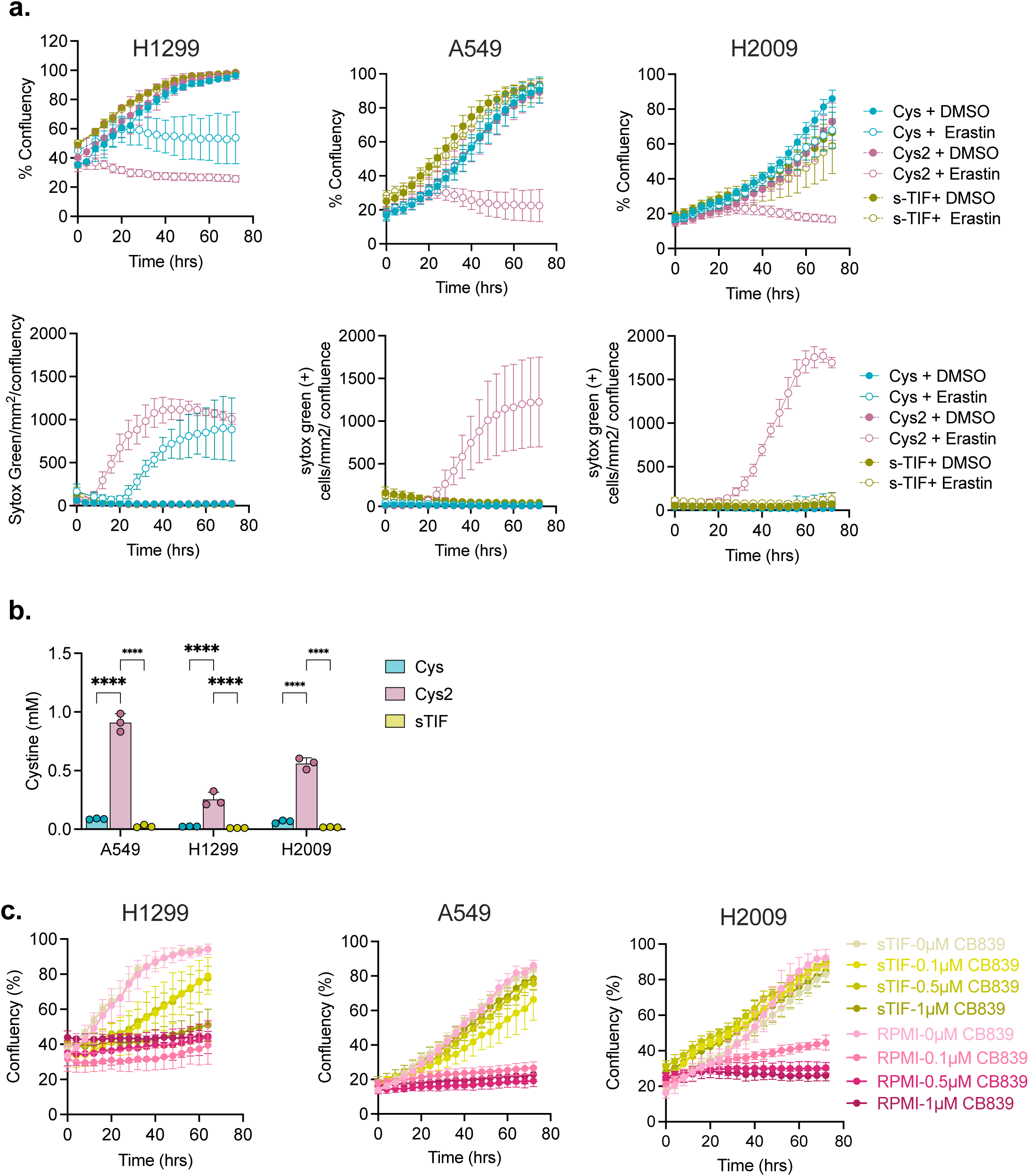
Lung sTIF supports lung cancer cell growth during metabolic and redox stress. **(a)** Live-cell imaging analysis of H1299, A549, and H2009 cells cultured in cysteine-containing medium (200 µM), cystine-containing medium (200 µM), or lung sTIF in the presence or absence of Erastin (5 µM for H1299; 10 µM for A549 and H2009). (Top) Percent confluence, (bottom) SYTOX Green-positive cells normalized to confluence over time (72 h).**(b)** Intracellular cystine concentration in A549, H1299 and H2009 cells cultured in cysteine (Cys: 200µM cysteine and 100µM TCEP), cystine (Cys_2_: 200µM cystine), and sTIF conditions for 8 h. **(c)** Live-cell imaging analysis of H1299, A549, and H2009 cells cultured in RPMI or lung sTIF and treated with the indicated concentrations of CB-839. Cell growth was monitored as percent confluence over time. All conditions contain 10% dialyzed FBS (dFBS). Data are mean ± s.d.; n = 4-6 biological replicates (**a,c**) or n = 3 biological replicates (**b**). Data are representative of 2 (**b**) or 3 (**a,c**) independent experiments.

**Supplementary figure 6:**
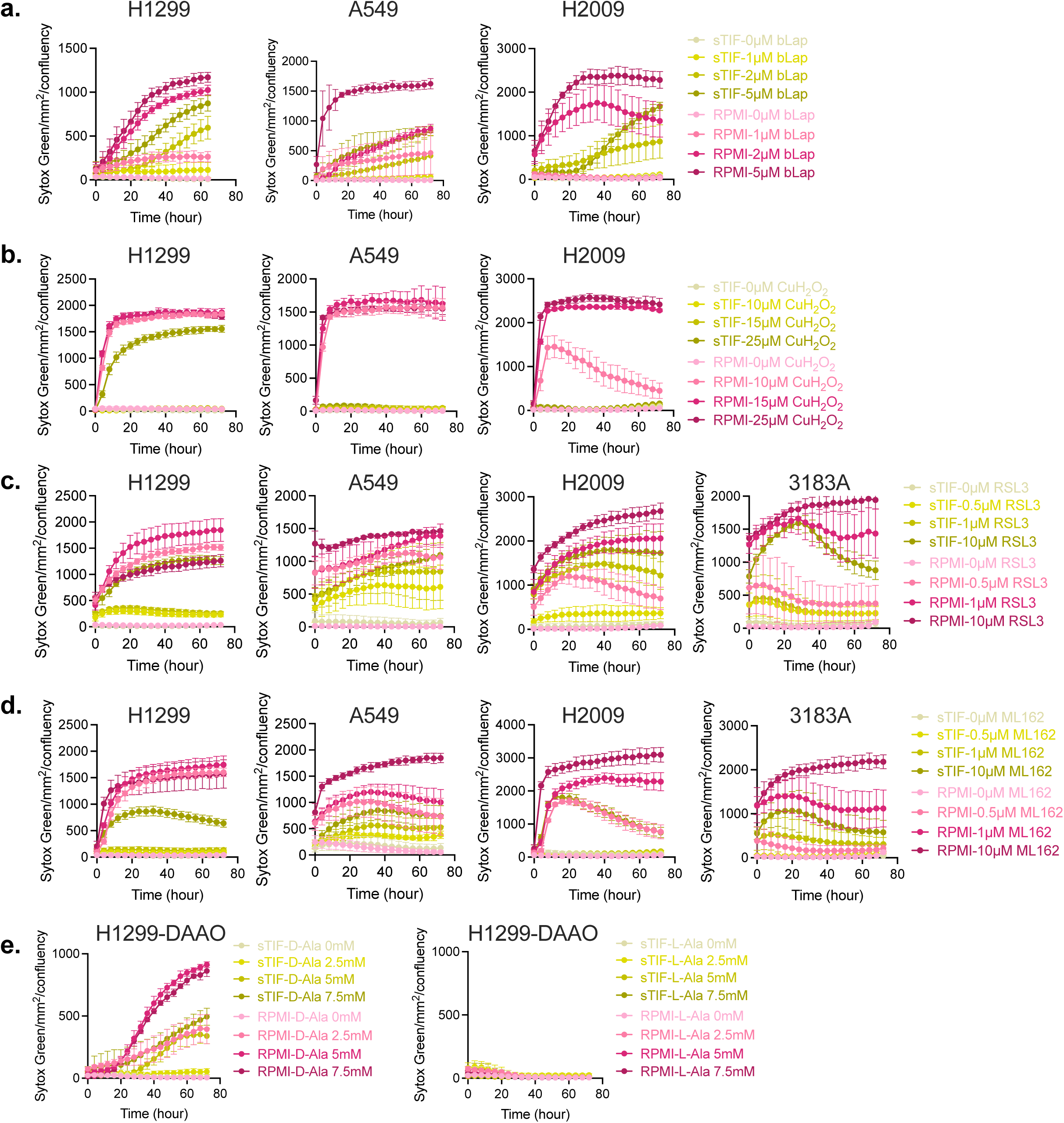
Lung sTIF alters lung cancer cell sensitivity to oxidative and ferroptotic stress. **(a-e)** SYTOX Green fluorescence signal normalized to confluence over time, in A549, H1299, H2009 and murine KP cells cultured at the indicated concentrations of β-lapachone (**a**), cumene hydroperoxide (CuH_2_O_2_)(**b**), RSL3 (**c**) or ML162 (**d**), and in D-AAO expressing H1299 cells cultured at the indicated concentration of D-Alanine or L-Alanine (**e**). Cells were cultured in RPMI or sTIF media +10% dFBS for 64-72 hours. D-AAO: D-Amino acid oxidase. Data are mean ± s.d.; n = 3-5 biological replicates. Data are representative of at least 2 independent experiments.

**Supplementary figure 7:**
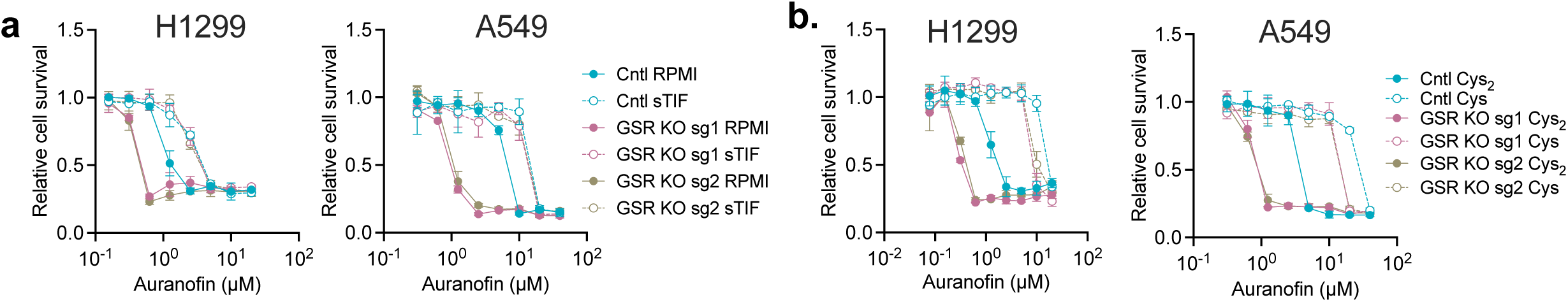
Lung sTIF alters lung cancer cell dependence on disulfide reductases. **(a-b)** Relative survival of control and GSR-knockout A549 and H1299 cells cultured in sTIF **(a)** or cysteine **(b)** conditions. (Cys: 200µM cysteine and 100µM TCEP), cystine (Cys_2_: 200µM cystine), and sTIF conditions and treated with the indicated concentrations of auranofin. All conditions contain 10% dialyzed FBS (dFBS). Data are mean ± s.d.; n = 3-5 biological replicates. Data are representative of at least 3 independent experiments.

